# Macropinocytosis mediates neurotropism of *Cryptococcus neoformans* in a human organoid model of the blood-brain barrier

**DOI:** 10.1101/2025.09.23.678106

**Authors:** Amelia B. Bennett, Dylan M. Lanser, Kiem Vu, Amita R. Sahoo, Matthias Buck, Angie Gelli

## Abstract

The opportunistic and neuroinvasive fungus, *Cryptococcus neoformans* (*Cn*), causes a life-threatening brain infection that despite treatment can cause long-term cognitive deficiencies. Studies have shown that *Cn* can infiltrate the central nervous system (CNS) through a transcellular route across the brain endothelium, however, the molecular process that drives brain endothelial cells to internalize *Cn* remains poorly defined. Here we examine the molecular interactions between fungal cells and the brain endothelium by utilizing a human 3D organoid model of the blood-brain barrier (BBB). We show that *Cn* exploits the process of macropinocytosis as the mechanism of endocytosis into brain endothelial cells by recruiting CD44 and EphA2 as a molecular complex. We identified two predicted binding sites on EphA2 for CD44, suggesting that the two structurally distinct regions may provide a molecular basis for cooperative signaling in brain endothelial cells that stimulate macropinocytosis as the mode of entry for *Cn*.

## Introduction

In a healthy brain, components of circulating blood that could damage the central nervous system (CNS) are excluded by the blood-brain barrier (BBB). The tightly connected endothelial cells, the low rate of vesicular transport and the highly-selective permeability of the BBB collectively ensure a protective environment for the CNS.^1^ Despite this, *Cryptococcus neoformans* (*Cn*), a neurotropic fungal pathogen and leading cause of meningoencephalitis in immunosuppressed adults^2^, has developed adaptations that promote its entry^3^ into- and proliferation within the CNS.^4,5^ Several lines of evidence support multiple routes of CNS invasion, including: (i) “Trojan Horse,” in which *Cn* infiltrates the BBB via infected phagocytes^6–12^, (ii) paracellular crossing of free *Cn* via the disruption of the BBB endothelial cell junction proteins^12–15^, and (iii) direct fungal interaction with brain endothelial cells leading to endocytosis and transcytosis of free *Cn*.^16–22^ The molecular mechanistic details of these routes, the propensity of *Cn* using one route versus another and whether all three routes are used during infection remains largely unresolved.

Utilization of endothelial cells and exploitation of endocytosis to cross the BBB via a transcellular path is central to *Cn*’s ability to infiltrate and survive within the CNS. Several *in vitro* and *in vivo* studies have demonstrated transcytosis as the primary mechanism driving *Cn* across the BBB.^3,16,23,24^ Evidence includes observational studies that found cryptococci moving freely through the bloodstream and crossing the brain endothelium independently of phagocytes^14^, and quantitative analyses that established a direct association between brain endothelial cells and *Cn in vivo*.^16,23^ Collectively, these data support recruitment of a transcellular pathway for BBB crossing, however the molecular process by which brain endothelial cells internalize *Cn* remains poorly understood.

One form of endocytosis is macropinocytosis, an actin-driven endocytic process that internalizes extracellular cargo via large vesicles referred to as macropinosomes.^25–27^ Based on previously observed indicators of macropinocytosis such as actin-supported membrane ruffles in brain endothelial cells infected with *Cn*^16,21^, actin disruptors preventing BBB crossing^13,21,28^ and the lack of evidence for other processes, we questioned whether *Cn* might exploit macropinocytosis as the mechanism of entry to the CNS.^29^

To investigate this, we utilized a human organoid model of the BBB. Multicellular BBB organoids have been used as a reliable and predictive platform that recapitulate features of the native BBB.^30^ The present study examines the molecular interaction between *Cn* and the brain endothelium. We previously found that *Cn* induced the non-canonical, ligand-free activation of the ephrin tyrosine kinase receptor, EphA2, in human brain endothelial cells.^21^ While we demonstrated that transcytosis of *Cn* across the BBB relied on the phosphorylation-dependent activity of EphA2^21^, the mechanistic details of the endocytic process remained unresolved. It was also unclear whether EphA2 played a role in recruiting CD44, the glycoprotein receptor for hyaluronic acid, and a known contributor to the transcellular pathway co-opted by *Cn* to cross the brain endothelium.^31–33^

## Results

### Cellular organization and validation of a human blood-brain barrier (BBB) organoid model of neural cryptococcosis

To examine whether macropinocytosis is the mechanism facilitating *Cn* internalization by the brain endothelium and transcytosis of *Cn* across the BBB, we constructed a 3D human organoid model of the BBB (Supp. Fig. 1a).^34,35^ To our knowledge, this is the first study that used a 3D BBB organoid model in the context of neurocryptococcosis. BBB organoids were generated by combining human immortalized brain endothelial cells (iBMECs) or primary brain endothelial cells, astrocytes and pericytes in a 1:1:1 ratio in commercially purchased low-adhesion 96-well plates specially designed to encourage the formation of unitary and uniformly sized (∼200 µm) organoids (Akura microplates, InSphero, Brunswick, ME, USA, Supp. Fig. 1b and inset). Approximately 1000 cells of each neurovascular cell type were combined to generate organoids, which formed spontaneously within 2-3 days and remained viable and static for at least 2 weeks. Scaffold support was not required as each cell type spontaneously self-arranged into a recapitulation of the cellular organization of the BBB.^34^

Immunofluorescence was used to examine expression of cellular markers of BBB organoids that were fixed and cryo-sectioned (Supp. Fig. 2). Surface-localized brain endothelial cells spontaneously established tight junctions, indicated by Claudin-5 expression, a transmembrane protein that is key to the physical establishment of the BBB through prevention of paracellular leakage (Supp. Fig. 2c, 2d, merge). Astrocytes and pericytes concentrated in the core of the organoids (Supp. Fig. 2e, 2f). Under these conditions, the size of the organoids remained unchanged over 14 days of culture, suggesting that endothelial cells appeared to regulate the proliferation of the pericytes and astrocytes encapsulated within the organoids. As the data supported the expected orientation of the three cell types in the organoids, we next examined the integrity of the endothelial barrier.

Using transmission electron microscopy (TEM), we assessed the integrity of the organoid endothelium and observed structures indicative of tight junctions between endothelial cells (Fig. 1a). The inability of the FITC-dextran (70 kDa) macromolecule to cross brain endothelial cells and infiltrate the organoid interior demonstrated BBB impermeability and confirmed an intact endothelial barrier encapsulating the organoid (Fig. 1c). In contrast, FITC alone, which is a much smaller molecule, freely crossed the endothelial barrier as expected (Fig. 1c). BBB organoids showed strong expression of Claudin-5 co-localized with the outer layers of the organoid, further confirming an intact barrier and also illustrating the precise location of the endothelial cell layer along the surface of the organoid (Fig. 1c). TEM analysis of BBB spheroids exposed to a high inoculum of fungal (*Cn*) cells revealed extensive plasma membrane ruffling confined to the endothelial cells that was in close proximity to *Cn* (Fig. 1d, 1e), consistent with cytoskeleton remodeling and in agreement with similar morphological changes observed in previous studies.^13,16,28^ BBB organoids consisting of either immortalized or primary human brain endothelia cells (Fig. 1f, 1g) were similar in organization and function.

**Fig 1.**
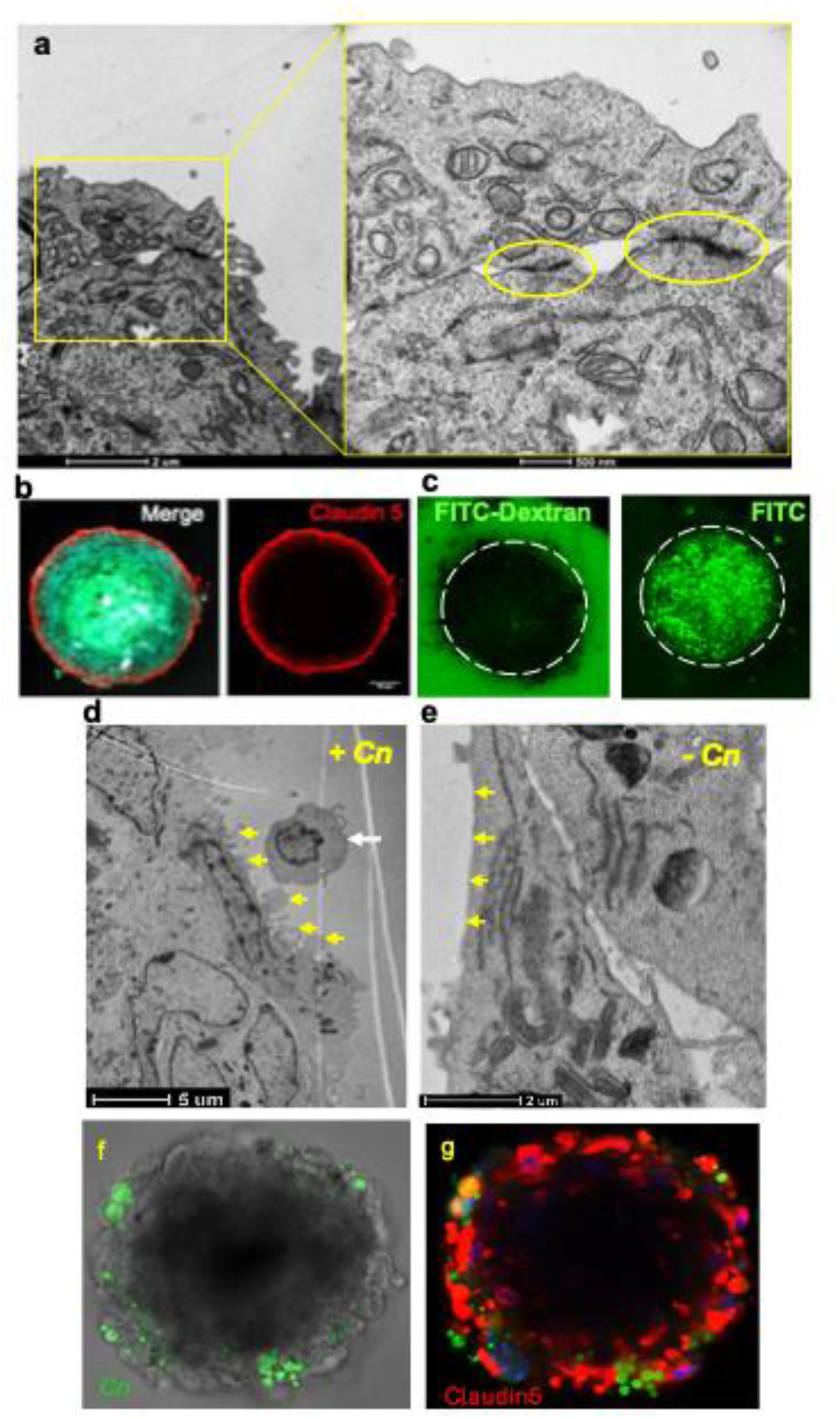
Human BBB organoids possess a functional endothelial barrier that is responsive to *Cn*. **a** TEM tight junctions between brain endothelial cells on the surface of an organoid. **b** Immunofluorescence image of the tight junction protein Claudin5 on the surface of fixed and sectioned BBB organoids, indicating the presence of tight junctions and position of endothelial cells. Interior of organoid stained with DAPI. **c** BBB organoids are not permeable to a large molecular weight dye (70 kDa FITC-Dextran, left panel), in contrast to small molecules like FITC (right panel). **d** TEM images of membrane ruffling and protrusions (yellow arrows) along the surface of brain endothelial cells exposed to *Cn* (white arrow), **e** in contrast to flat endothelial cell surface (yellow arrows) on BBB organoids not exposed to *Cn*. **f, g** Immunofluorescence image of BBB organoids consisting of primary human brain endothelial cells express Claudin-5 (red) and show infiltration of *Cn* (green) indicating similar morphology and function to organoids with immortalized brain endothelial cells.

### Treatment of the BBB with an inhibitor of macropinocytosis prevented *Cn* crossing of the BBB

Next, we assessed whether the BBB organoids permitted *Cn* crossing to a degree that could be robustly analyzed. *Cn*, labeled in green with an antibody to the GXM of the polysaccharide capsule, appeared to penetrate the Claudin-5-labeled endothelial layer of BBB organoids (Fig. 2a-c). Numerous *Cn* were observed within endothelial cells and near DAPI-stained astrocytes or pericytes that were closest to the endothelial layer, but appeared to be largely excluded from the astrocyte core (Fig. 2a (inset), 2b). We also observed sloughing off of *Cn* capsule, indicated by green puncta (Fig. 2c), consistent with earlier studies that found remnants of the polysaccharide capsule with reactive astrocytes adjacent to cryptococcomas during infection.^36–38^ We found that *Cn* was readily internalized by BBB organoids, in contrast to the sibling species, *Cryptococcus deuterogattii* (*Cd*), and to the negative control, *Saccharomyces cerevisiae* (*Sc*) (Fig. 2d, 2e). Collectively, these results suggest that the BBB organoids are highly functional entities able to distinguish neuroinvasive pathogens from others, and could serve as a suitable model to examine the mode of *Cn* entry.

**Fig 2.**
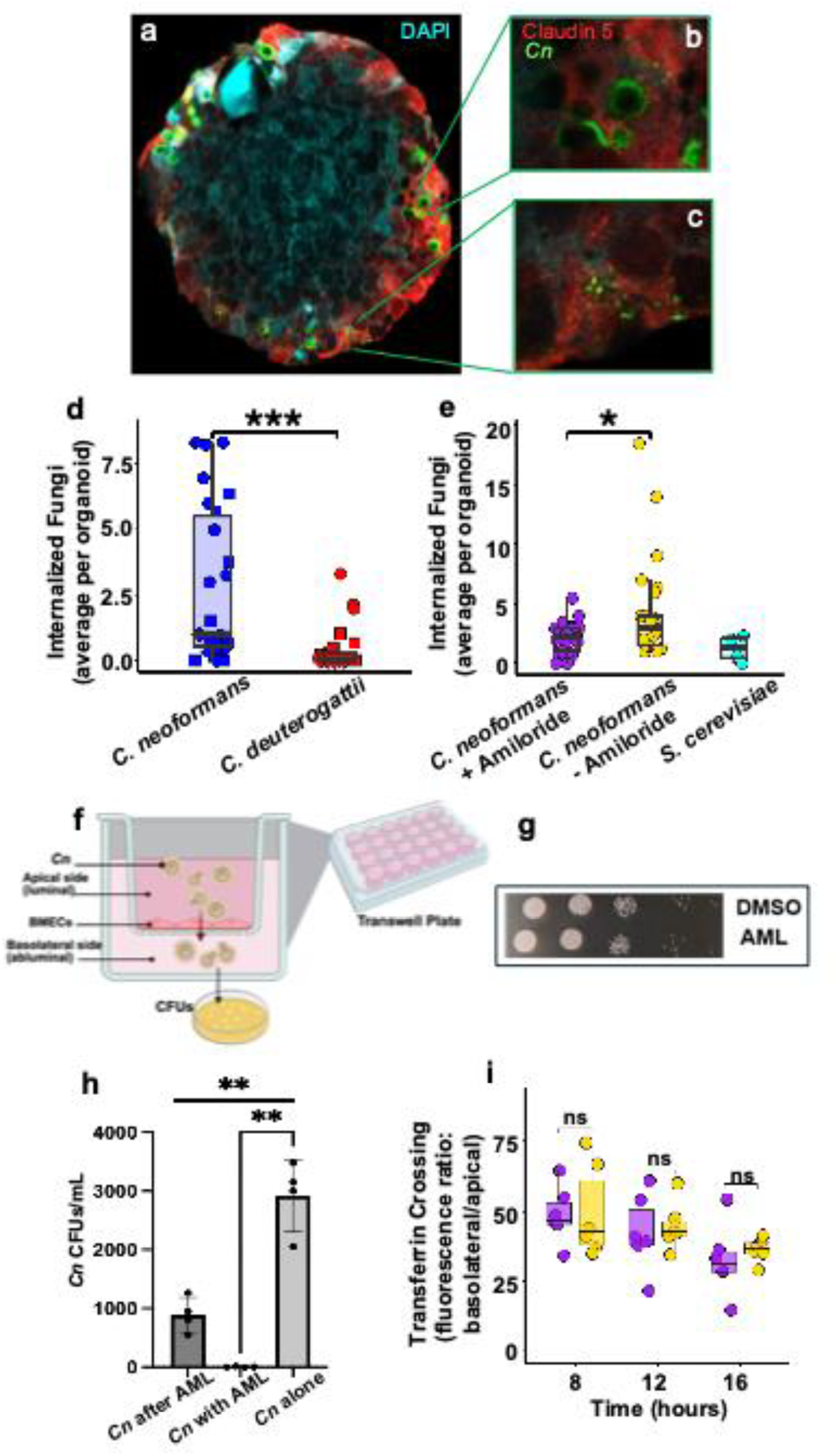
Treatment of BBB organoids with Amiloride, a potent inhibitor of macropinocytosis, prevents fungal cells from crossing the endothelial barrier of BBB organoids. **a** Fluorescent image of BBB organoid incubated with *Cn* (green) for 72 h, fixed, sectioned and imaged on a confocal microscope. *Cn* stained with anti-capsule antibody (anti-GXM 18b7, 1:300) kindly provided by A. Casadevall. Claudin-5 expression shown in red. **b** Enlarged image shows fungal cell (green) internalized by Claudin-5 (red)-expressing endothelial cells. **c** Degraded and/or released *Cn* capsule components shown in green. d *Cn* invades BBB organoids at a significantly higher rate than *Cd* at 48 h (two-sample t-test with Welch’s correction: t = 4.01, df = 28.59, *p* < 0.0004). **e** Amiloride (250 µM) treatment significantly reduces (t = −2.46, df = 28.23, *p* = 0.020) *Cn* internalization to a level comparable to non-pathogenic *Saccharomyces cerevisiae* (t = 1.53, df = 4.50, *p* = 0.194). Each data point represents the average number of internalized fungal cells per BBB organoid across multiple images of fixed and cryo-sectioned organoids. For (d) and (e), each data point represents the average number of internalized fungal cells per BBB organoid across multiple images of fixed and cryosectioned organoids. **f** Schematic of the transwell human BBB model. To assay for BBB crossing, fungal cells or fluorescent macromolecules were first added to the upper chamber (luminal side) and then collected from the bottom well (abluminal side) for either CFU enumeration or fluorescence quantification. **g** Spot assay comparing growth of *Cn* on minimal media following incubation for 24 h with 250 µM amiloride or DMSO control in endothelial media at 37°C, 5% CO_2_. **h** Treatment of iBMECs with amiloride (250 µM) inhibits *Cn* crossing in an *in vitro* BBB transwell model. iBMECs were either pre-treated with amiloride or treated with amiloride along with *Cn*. Statistical analysis of the transwell assay was done using two-tailed unpaired t-test with Welches correction (t = 9.504, df = 3.00, \*\**p* = 0.0025) or one-way ANOVA (GraphPad Prism 10 software). **i** In the transwell model, the presence of amiloride (purple) makes no significant difference in the rate of transferrin transcytosis relative to the vehicle control (yellow). The relative fluorescence ratio (RFU) was measured at 8 h, 12 h, and 16 h post-treatment. Non-significant differences were determined by Welch two-sample t-test between treatment and control at 8 h (t = −0.12, df = 8.27, *p* = 0.91), 12 h (t = −0.34, df = 8.42, *p* = 0.74), and 16 h (t = −0.65, df = 6.13, *p* = 0.54) (R v4.2.2).

Macropinocytosis is uniquely susceptible to amiloride (AML), a chemical inhibitor of Na^+^/H^+^ exchange that inhibits macropinocytosis by lowering endomembrane pH and thereby inhibiting actin polymerization.^39^ Treating BBB organoids with AML prior to *Cn* exposure significantly reduced the number of fungal cells internalized, in contrast to the vehicle treated control (Fig. 2e; Supp. Fig. 3). Fungal cell viability was not impacted at the concentration of AML used in these assays (Fig. 2g). To confirm this result and by extension the BBB organoid-based model, we used a well-established 2D transwell-based model of the BBB^40,41^ to investigate *Cn* crossing of AML-treated cultured human brain microvascular endothelial cells (HBMECs)^41^ (Fig. 2f). In this model, HBMECs (similar to the cells used in the organoids), were grown on a transwell resulting in an intact, impermeable endothelial barrier that separated the bottom well (abluminal/basolateral) from the upper chamber (luminal/apical side).^21,40,42^ AML treatment was administered to the top chamber either before exposure to *Cn* or added to the fungal cell inoculum. The number of fungal cells crossing the endothelial barrier was reduced by ∼3-fold with AML compared to the vehicle treatment, and the data were consistent with the results from the BBB organoids (Fig. 2h). Employing the same 2D transwell model of the BBB, the crossing of transferrin - used here as a previously validated negative control for macropinocytosis-dependent transcellular crossing^43^ - was examined 8 h, 12 h and 16 h post treatment with AML. The relative fluorescence unit (RFU) ratios of transferrin in the upper chamber versus the bottom well were not significantly different at all time points, suggesting that clathrin-mediated endocytosis of transferrin was not inhibited by AML (Fig. 2i). The AML-mediated inhibition of *Cn* internalization and crossing in both 2D and 3D *in vitro* BBB models was strongly indicative of macropinocytosis involvement.

### Brain microvascular endothelial cells challenged with *Cn* induced the internalization of a macromolecule, further implicating macropinocytosis

Serum starvation of cells can induce macropinocytosis thereby enabling cells to engulf extracellular cargo including macromolecules.^44,45^ To visualize the process of macropinocytosis in brain endothelial cells exposed to *Cn,* we examined the fate of a 70 kDa FITC-labeled dextran macromolecule (Fig. 3).^46^ Brain endothelial cells were either starved of serum or fed with medium containing serum (non-serum starved) for 24 h and subsequently exposed to *Cn* for 1 h in the presence of FITC-dextran. We observed a *Cn*-induced uptake of dextran in cells that was independent of serum starvation (Fig. 3d). Distinct peri-nuclear puncta (green), indicative of dextran uptake, observed in serum starved (Fig. 3a) and *Cn*-treated (Fig. 3b, 3d) were found in close proximity to cryptococci (red) (Fig. 3d). Quantification of the internalized dextran signal suggested that *Cn* induced significant uptake of the dextran macromolecule in non-serum starved cells to levels comparable to serum starvation in cells not exposed to *Cn* (Fig. 3e). Moreover, the uptake of FITC-dextran induced by *Cn* in non-serum starved cells was significantly diminished by AML-treatment, further implicating macropinocytosis (Fig. 3f).

**Fig 3.**
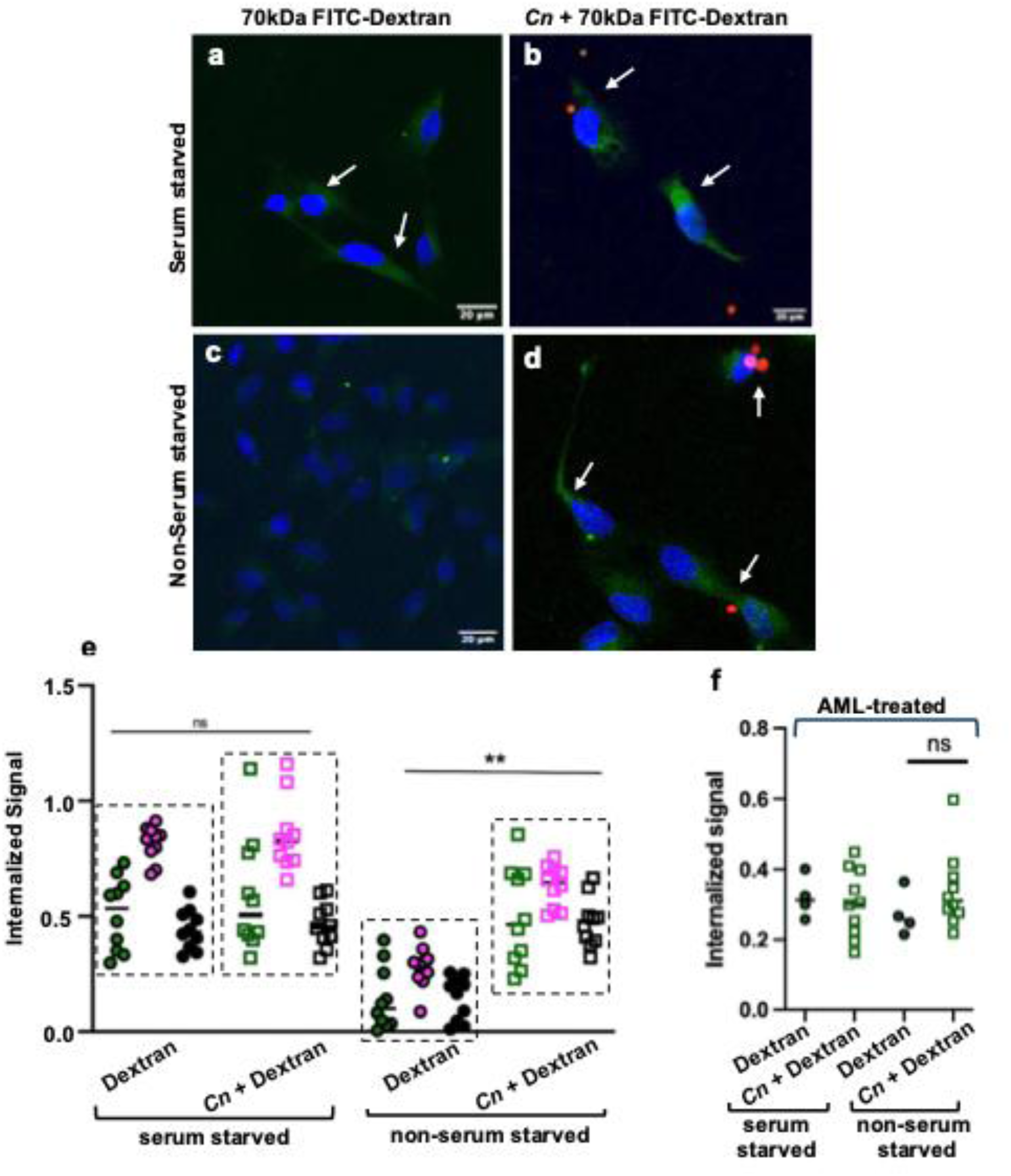
***Cn* induces the internalization of the macromolecule, 70kDa FITC-Dextran, by brain endothelial cells independent of serum starvation. a – d** Exposure of non-serum starved human iBMECs (stained with DAPI, blue) to *Cn* promotes internalization of dextran indicated by perinuclear staining of dextran (green puncta) that co-localizes with fungal cells (*Cn,* shown in red). This is in contrast to non-serum starved iBMECs not treated with *Cn* (negative control). Serum starvation of iBMECs (positive control) leads to dextran uptake. iBMECs were serum starved for 24 h and then exposed to 0.5mg/mL FITC-Dextran (70 kDa) and treated with *Cn* (MOI 10) for 1 h. iBMECs were washed of extracellular fluorescent signal, fixed and analyzed via confocal microscopy. **e** Internalized FITC-dextran signal was quantified by determining the internalized signal of FITC-Dextran within the area of each cell (internalization index) using ImageJ.^46^ Three biological replicates shown for each treatment. Each data point represents the average internalization index within one microscopic image. Statistical analysis was done using a two tailed, unpaired t test (n = 10, per group, t= 5.934, df=18, \*\**p* = 0.007) (GraphPad Prism 10 software). **f** Amiloride inhibits FITC-dextran uptake by iBMECs. iBMECs serum starved and non-serum starved were treated with 250 µM amiloride for 1 h prior to exposure to *Cn* (MOI of 10) and 0.5 mg/mL FITC-dextran. Internalized FITC-Dextran signal was quantified by determining the internalized signal of FITC-Dextran within the area of each cell (internalization index) using Image J. Each point represents the average internalization index within one microscopic image (n > 3). Statistical analysis was done using a two tailed, unpaired t test with Welch correction (t= 1.412, df =9.5, *p* = 0.1897) (GraphPad Prism 10 software).

### *Cn* crossing of the BBB in the organoid model was contingent on EphA2 activity

We next examined the contribution of the ephrin receptor tyrosine kinase, EphA2, to the neuroinvasion of *Cn* in BBB organoids that were genetically modified with the CRISPR/Cas9 system to knock out EphA2 in brain endothelial cells. The EphA2 knockout (KO) cells expressing Red Fluorescence Protein (RFP) exhibited the same peripheral localization as wild type (WT) endothelial cells (Fig. 4a). Western blot analysis of EphA2 KO cells confirmed the lack of EphA2 protein expression (Fig. 4b).

**Fig 4.**
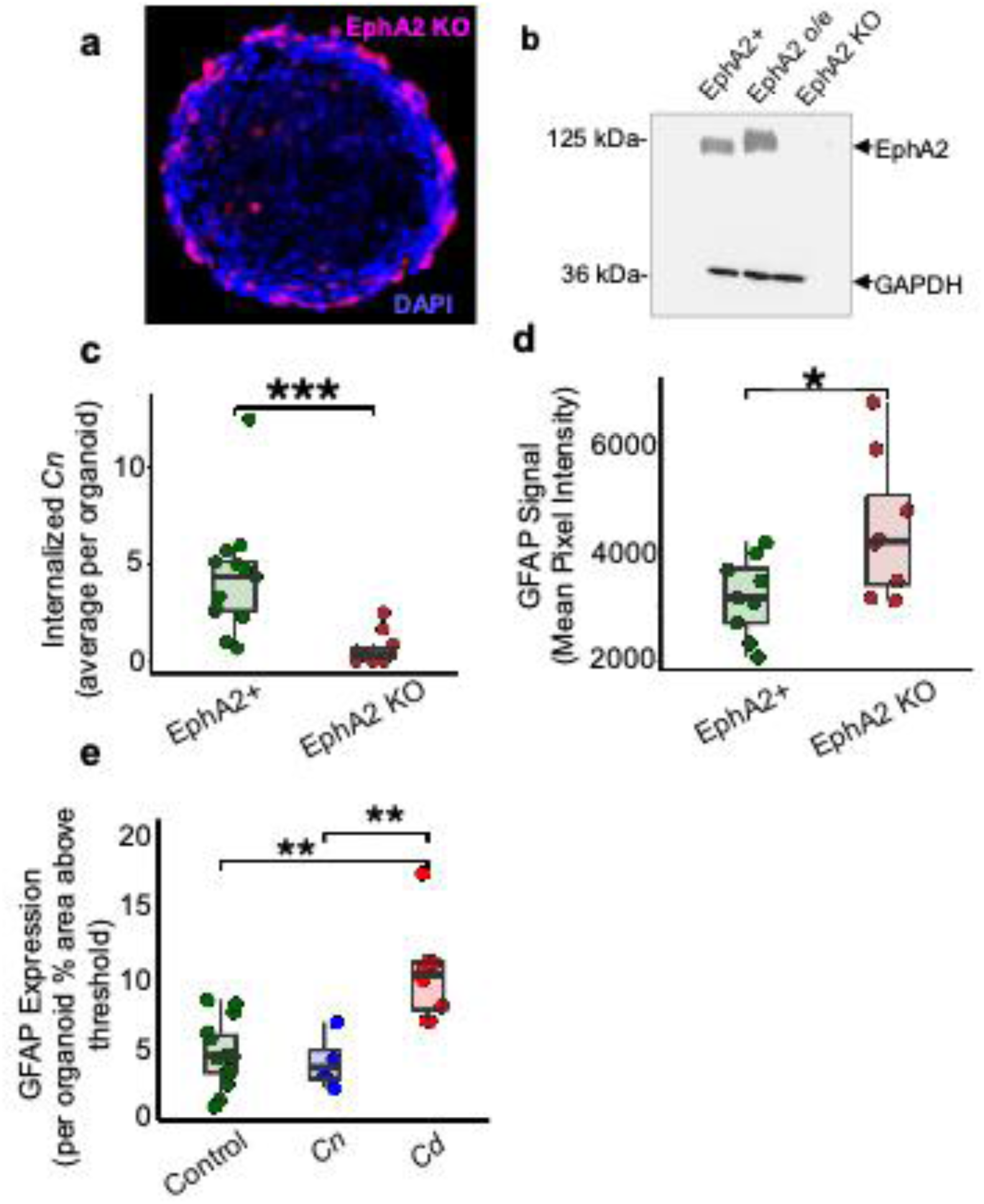
EphA2 expression promotes barrier integrity while facilitating *Cn* Internalization. **a** Representative fluorescent image of BBB organoids formed from EphA2 knockout (KO) human brain endothelial cells. EphA2 KO cells express RFP (magenta) against DAPI (blue) counterstaining. **b** Western blot analysis (1:1000 dilution of rabbit anti-Human EphA2; Cell signaling technology Inc. D4A2) (1:10,000 dilution of Goat Anti-Rabbit IgG H&L HRPab6721; Abcam Inc., Cambridge, MA, USA) confirming loss of EphA2 protein expression in EphA2 KO organoids, in contrast to controls (EphA2+ and EphA2 over-expression (o/e)). **c** Internalization of *Cn* (*Cryptococcus neoformans*) by BBB organoids is dependent on EphA2 expression. Organoids (EphA2+ or EphA2 KO brain endothelial cells) were exposed to CFSE-stained *Cn* for 48 h, after which organoids were sectioned and analyzed for *Cn* invasion (Welch two-sample t-test: t = 4.28, df = 13.94, *p* < 0.0008)(R v.4.4.2). **d** Comparison of GFAP expression between BBB organoids formed with EphA2 KO versus EphA2+ brain endothelial cells. Organoids deficient in EphA2 show significantly higher levels of GFAP expression compared to EphA2+ organoids. Quantification was performed on GFAP-antibody probed organoid sections by applying a uniform threshold to images and measuring the percent area in the relevant channel above the threshold (Welch two-sample t-test: t = 2.42, df = 10.68, *p* = 0.0343)(R v.4.2.2). **e** Organoids exposed to *Cd* (*Cryptococcus deuterogattii*) show increased GFAP expression in stark contrast to organoids exposed to *Cn* after 48 h of exposure (Welch two-sample t-test comparing *Cn* & *Cd* treatments: t = 3.84, df = 9.38, *p* = 0.003, comparing *Cd* & Control treatments: t = 4.25, df = 10.57, *p* = 0.0015)(R v.4.4.2). For panels c – e, each data point represents the average value for an organoid across several images.

Organoids constructed with EphA2 KO endothelial cells were exposed to *Cn* for 24 h, after which the number of endocytosed fungal cells was quantified (Fig. 4c). Significantly fewer fungal cells were internalized by the organoids deficient in EphA2. Conversely, over-expression of EphA2 resulted in an increased rate of *Cn* crossing (Supp. Fig. 4). These results were consistent with our prior study using the 2D BBB transwell model showing that EphA2 is required for *Cn* to cross the brain endothelium.^21^ This also further demonstrated that the lack of EphA2 may not have compromised the integrity of the organoid endothelium to the extent that it permitted *Cn* crossing.

To assess whether organoid astrocytes were affected by the absence of EphA2 in endothelial cells, GFAP, a marker of astrocyte activation and indication of neuroinflammation, was quantified in intact organoids. The intensity of GFAP expression was significantly higher in organoids deficient in EphA2 when compared to wild type (EphA2+) organoids (Fig. 4d). Notably, GFAP expression increased significantly when organoids were exposed to *C. deuterogattii* (*Cd*) a fungal species known to cause disease primarily in immunocompetent individuals.^47,48^ This result was contrary to the levels of GFAP expression in organoids challenged with *Cn*, which were similar to control (Fig. 4e). These data are consistent with numerous studies that have reported a milder neuroinflammatory response associated with *Cn* brain burden compared to other neurotrophic pathogens, and further underscore the ability of BBB organoids to discern differences that the more commonly used 2D transwell BBB models overlook.^49,50^

### *Cn-*induced activity of GTP-bound Cdc42 is dependent on both EphA2 and CD44

Macropinocytosis, initiated by actin-polymerization and plasma membrane folding, produces cell surface ruffles that encapsulate extracellular cargo into macropinosomes. The entire process is regulated by signaling pathways, including small Rho GTPases such as Cdc42.^51^ To test the hypothesis that EphA2 promotes macropinocytosis of fungal cells by influencing actin remodeling through downstream signaling of Cdc42, we employed an activation assay to examine Cdc42 activity. GTP-bound (active) Cdc42 was detected in brain endothelial cells exposed to *Cn* with high levels of Cdc42 activity relative to unexposed cells (Fig. 5a). EphA2 KO endothelial cells that had been exposed to the same *Cn* inoculum had significantly reduced GTP-bound Cdc42 (Fig. 5a), suggesting that *Cn* stimulation of active Cdc42 was dependent on EphA2 activity. We confirmed by western blot analysis that Cdc42 protein expression levels were comparable in all endothelial cell lysates (Fig. 5c).

**Fig 5.**
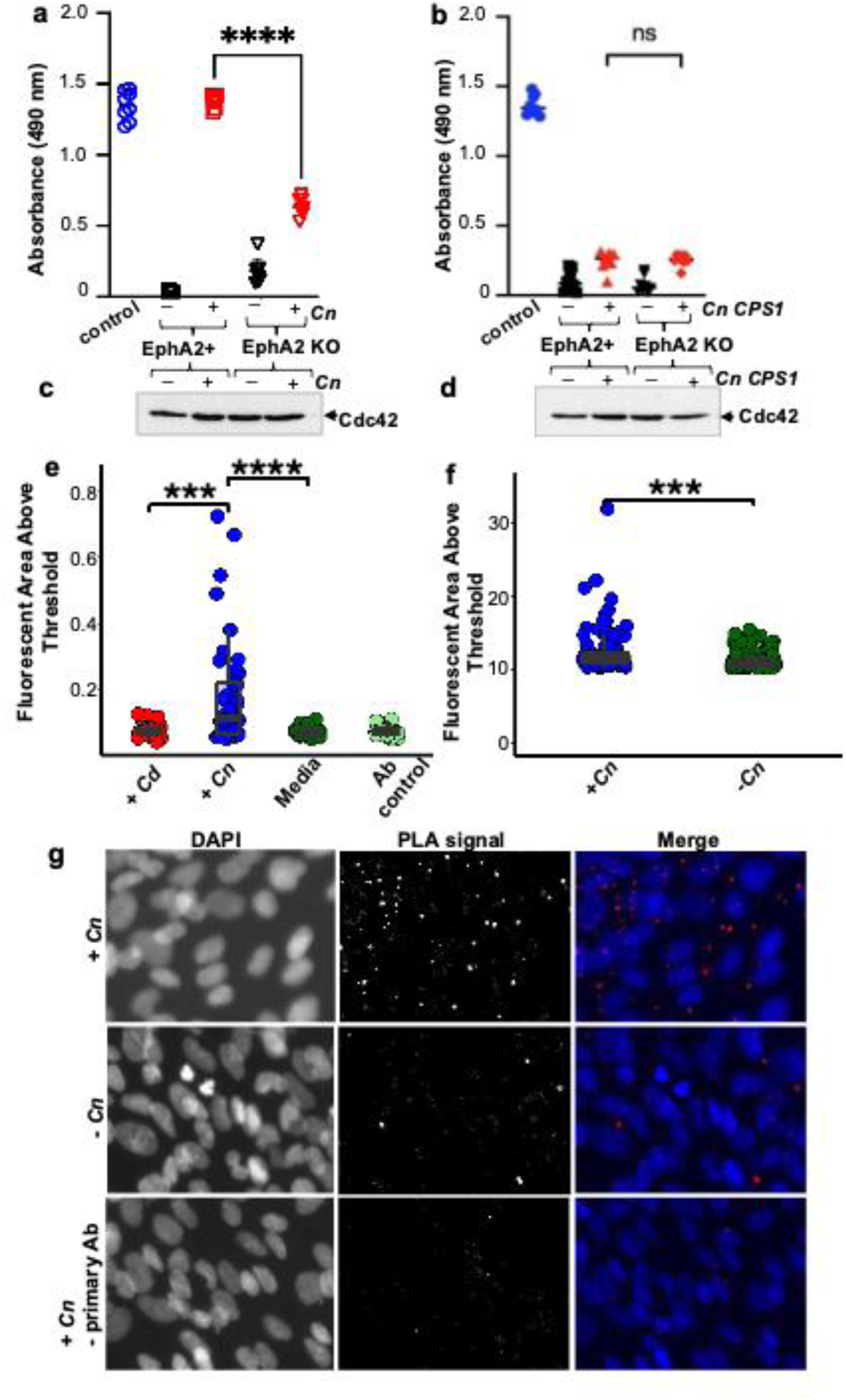
*Cn* activates GTP-bound Cdc42 via recruitment of a CD44-EphA2 molecular complex. **a** *Cn* induced stimulation of GTP-bound Cdc42 is contingent on EphA2 expression and **b** CD44 engagement. EphA2+ and EphA2 KO human brain endothelial cells were treated with either **a** *Cn* or **b** *cps1Δ Cn* deletion mutant at an MOI of 10 for 30 min and then lysed. Untreated and treated cell lysates were analyzed for GTP bound Cdc42 (active Cdc42) using the Cytoskeleton G-LISA assay. Data are expressed as a mean ± SD of three independent experiments. Statistical analysis was done using a two tailed unpaired t-test (n = 8, t = 0.2156, df = 14, \*\*\*\**p* < 0.0001)(GraphPad Prism 10 software). **c, d** Expression of Cdc42 protein in lysates was confirmed by western blot analysis of nitrocellulose membrane probed with primary antibody to amino acid 150-182 of Cdc42 protein. (1:800 dilution of mouse anti-human Cdc42; cytoskeleton Inc. ACD04) (1:10,000 dilution of Goat anti-Mouse IgG H&L HRP ab6789; Abcam Inc., Cambridge, MA, USA). **e, f** *Cn* recruits a CD44-EphA2 protein complex in brain endothelial cells, detected by Proximity Labeling Assays (PLA - DuoLink) of EphA2 and CD44 in **e** mouse primary brain endothelial cells and **f** human brain endothelial cells (EphA2+, iBMEC). The *in-situ* interaction was quantified as the area in an image above a universally applied threshold. Significantly higher fluorescence was observed in *Cn* (+) versus *Cn* (-) treatments in both mouse primary cells (t = 4.40, df = 43.13, *p* < 0.0001) and human cells (t = 3.63, df = 131.88, *p* = 0.0004). In mouse primary cells, there was significantly higher signal in the *Cn* (+) compared to the *Cd* (+) treatments (t = 4.08, df = 43.63, *p* < 0.0002)(R v.4.4.2). Mouse primary brain endothelial cells were exposed to *Cd*, *Cn* media or a no primary antibody control. **g** Representative fluorescent images of PLA in human brain endothelial cells (EphA2+, iBMECs). Top row: iBMECs exposed to *Cn* for 90 min. Middle row: control panel iBMECs not exposed to *Cn*. Bottom row: control treatment in which iBMECs were exposed to *Cn* but omitting primary antibodies (anti-CD44 and anti-EphA2) Left column: Nuclear stain. Middle column: Fluorescent puncta (PLA-red probe) are indicative of target proteins (EphA2 and CD44) in proximity (within 40 nm) to each other. Right column: merged images.

Next, we sought to examine whether adherence of *Cn* to brain endothelial cells via CD44 was required for the EphA2-mediated macropincytosis of *Cn*. Previous studies found that a hyaluronic acid-deficient strain of *Cn*, which lacks the gene encoding hyaluronic synthase (*CPS1*), failed to bind CD44, a transmembrane glycoprotein receptor for hyaluronic acid, and was defective at crossing the BBB.^31–33^ Employing the same Cdc42 activation assay as performed above, we detected significantly reduced levels of active GTP-bound Cdc42 in wild type (EphA2+) and EphA2 KO brain endothelial cells exposed to the *CPS1* deletion strain of *Cn* (Fig. 5b). The lack of activity was not due to reduced or absent expression of Cdc42 in cell lysates as evidenced by western blot analysis (Fig. 5d). Collectively, the data suggest that *Cn* engages both CD44 and EphA2 to promote Cdc42-mediated signaling and thereby induce macropinocytosis.

### *Cn* recruits CD44 and EphA2 as a complex in brain endothelial cells

The dependence of *Cn* on both EphA2 and CD44 to adapt macropinocytosis and cross the BBB, led us to question whether EphA2 could form a molecular complex with CD44 in brain endothelial cells. To probe this possibility, we employed AlphaFold3 (AF3) ^52^ to predict potential interaction interfaces between the EphA2 extracellular region and CD44. The structural predictions suggested two plausible binding sites for CD44: one located in the EphA2 ligand-binding domain (LBD) and a second in the fibronectin domains (FN1–FN2) positioned proximal to the plasma membrane (Fig. 6a). The predicted models yielded ptm scores of 0.49 and iptm scores of 0.13/ 0.11, indicating relatively low confidence in the inter-protein interface. Such low iptm values are not unexpected, as no structural evidence for EphA2–CD44 interactions currently exists in structural databases, making it challenging for AF3 to accurately capture the interface. Nevertheless, AlphaFold models can still serve as useful hypotheses for exploring possible binding modes. To test whether the predicted complexes could form stable assemblies, we carried out molecular dynamics (MD) simulations on the LBD:CD44 and FN:CD44 models independently. As a positive control, we simulated the EphA2 LBD bound to ephrinA5, using the crystal structure as the starting model.^53^ In all cases, the complexes remained stable throughout 200 ns of simulation in solution. The backbone RMSD profiles (Supplementary Fig. 6. a–c) exhibited minimal fluctuations, indicating that both CD44-binding modes are structurally compatible with EphA2 and can persist on the relevant timescale. We next clustered the simulation trajectories, using an RMSD cutoff of 5 Å for LBD:CD44 and FN:CD44 complexes, and 2 Å for the LBD:ephrinA5 control system. The dominant conformation of the LBD:CD44 complex (Fig. 6b) highlighted specific EphA2 residues, including Q56, Y65, D61, and F156, that formed stable contacts with CD44. Interestingly, the same EphA2 residue Q56 also mediates interactions with ephrinA5, together with R103, D53, and G39, in the LBD:ephrinA5 complex (Supplementary Fig. 6d). This overlap suggests that CD44 and ephrinA5 may compete for binding to the same functional interface of EphA2. By contrast, the dominant FN:CD44 conformation (Fig. 6c) revealed interactions involving both FN1 and FN2 domains, suggesting that CD44 can also engage EphA2 through a distinct, membrane-proximal site. To gain deeper mechanistic insights, we analyzed intermolecular contacts over the course of the simulations (Supplementary Fig. 7a-c). The interaction analysis confirmed that CD44 engages the LBD of EphA2 at an interface overlapping with ephrinA5 binding, while also being capable of forming a secondary, non-overlapping interaction with the FN domains. The ability of CD44 to bind EphA2 at two structurally distinct regions may provide a molecular basis for cooperative signaling in endothelial cells. In particular, competition with ephrin ligands at the LBD could modulate canonical EphA2 signaling, whereas FN-mediated interactions may stabilize CD44 association near the membrane to facilitate macropinocytic uptake.

**Fig 6.**
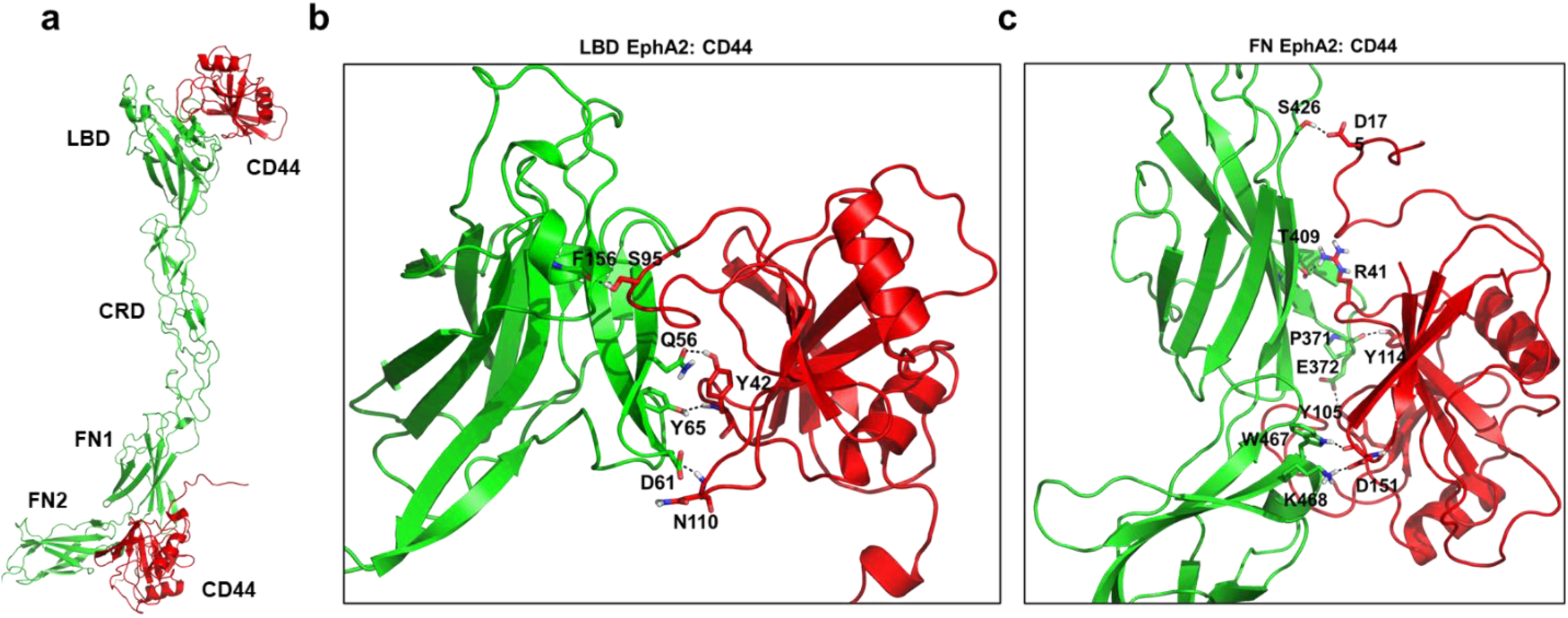
**Alpha-Fold-Multimer structure prediction indicates reliable binding between EphA2 and CD44**. **a** Cartoon representation of the AlphaFold3-predicted model of the EphA2 structure. extracellular region in complex with CD44, highlighting two distinct putative binding interfaces: the ligand-binding domain (LBD) (ipTM = 0.13; pTM = 0.49) and the fibronectin domains (FN1-FN2) (ipTM = 0.11; pTM = 0.49). **b-c** Representative binding conformations of the LBD: CD44 and FN: CD44 complexes obtained from molecular dynamics simulations. Shown here are the major conformational clusters derived from all replica trajectories. EphA2 is depicted in green and CD44 in red cartoon representations. For clarity, only the principle interfacial hydrogen bonds are displayed.

To visualize the CD44 and EphA2 protein-protein complex, we used an *in-situ* fluorescence-based proximity ligation assay (PLA, DuoLink) in both primary and immortalized brain endothelial cells challenged with *Cn.* The detected fluorescent signal, evidenced by the red puncta in brain endothelial cells challenged with *Cn*, were indicative of a molecular interaction between EphA2 and CD44 (Fig. 5g, 5h). Quantification of the fluorescence signal detected in mouse primary brain endothelial cells (Fig. 5e) and in immortalized human brain endothelial cells (Fig. 5f) revealed stimulation of the EphA2-CD44 protein interaction by *Cn* in both cell types, which was not observed in brain endothelial cells exposed to *Cd*.

## Discussion

A key innovation of the current study is the use of the 3D human BBB organoid model for the first time to elucidate mechanistic details by which *Cn* engages and crosses the BBB. A crucial feature of the organoid, vital to replicating the functionality of the BBB, is the direct contact of endothelial cells, pericytes and astrocytes, which collectively maintain the barrier integrity.^30^ Despite some limitations, human BBB organoids are a highly versatile and translational tool that overcome many of the constraints posed by 2D *in vitro* BBB models.^30^ By leveraging human BBB organoids in our studies, we determined that *Cd* and *Cn* elicit distinct responses from endothelial cells and astrocytes, which would have been difficult to discern in other BBB models. We also demonstrated that the BBB organoids are amenable to genetic manipulation. We engineered BBB organoids with endothelial cells that lacked EphA2 and detected a concomitant increase in GFAP+ astrocytes that likely acted in a compensatory manner to reinforce the barrier, as evidenced by the inability of *Cn* to cross the BBB.^54,55^ These results provide mechanistic insight into *in vivo* studies that found EphA2 deficiency preserved BBB integrity.^56,57^

In the present study we have made several compelling observations of the endocytic mechanism responsible for internalizing *Cn* and driving transcytosis across the brain endothelium. Despite the challenging nature of demonstrating the engulfment and transport of a pathogen by macropinocytosis, due to the non-specificity of this endocytic pathway, the data presented here strongly implicate macropinocytosis as the mechanism of entry. Exploitation of this process by *Cn* is of particular interest because the paucity of fluid phase endocytosis (i.e., macropinocytosis) is a distinguishing feature of the brain endothelium.^58–60^ At the same time, other pathogens have also adapted this mode of entry into non-phagocytic cells,^61–68^ which may have evolved from interactions either with innate immune cells that utilize macropinocytosis to survey and internalize the bulk extracellular milieu, or from single-celled eukaryotes that use macropinocytosis as a feeding-mechanism.^69^ Unlike other endocytic processes, macropinocytosis^70–73^ is not triggered by cargo binding or direct contact with large particles. Moreover, the larger size of macropinosomes (0.2-10 μm in diameter) would be the appropriate size for *Cn* (4-6 μm), unlike canonical endocytic pathways that typically produce vesicles less than 0.1 μm in diameter.^26,69^ While macropinocytosis is considered to be essentially nonselective (receptor-independent), recent evidence suggest that ligand-bound integrins and receptor tyrosine kinases may provide some degree of selectivity to this process.^74^

In a previous study, we demonstrated that transcytosis of *Cn* is mediated by a *Cn*-induced, ligand-independent activation of EphA2, potentially initiated by CD44, but the underlying mechanism had not been determined.^21^ Moreover, no studies have linked EphA2 and CD44 in the context of fungal neuroinvasion, although multiple lines of evidence have implicated their individual roles.^18,21,32,33^ In the present study, we found that CD44 and EphA2 are both critical for the successful entry and transcytosis of free *Cn.* Collectively our data support a model in which *Cn* has adapted macropinocytosis as the mechanism of entry to the CNS by recruiting both EphA2 and CD44 (Fig. 7). This is based on two key observations: (i) we detected a molecular interaction between EphA2 and CD44 in both human and primary mouse brain endothelial cells that is stimulated by *Cn*; and (ii) the absence of EphA2 or a failure to engage CD44 via hyaluronic acid-binding prevented the downstream activation of Cdc42, a key activator of macropinocytosis.^75,76^ Our model is further supported by the established independent roles of CD44 and EphA2 in *Cn* adherence to- and crossing of the brain endothelium, both of which require cytoskeleton remodeling, and activation of Rho-GTPase signaling cascades - activities attributed to both CD44 and EphA2.^77^ Our model suggests that the *Cn*-induced phosphorylation of EphA2 we previously reported as a required event for transcytosis^21^, is mediated by the association of EphA2 with CD44 upon binding *Cn*. This transactivation of EphA2 may involve PKC, a serine threonine kinase required for transcytosis of *Cn*^18^ and known to phosphorylate EphA2 independent of ligand binding (non-canonical signaling).^78,79^ The culmination of EphA2-CD44 complex formation would activate Cdc42-mediated signaling, and thereby trigger macropinocytosis. Our *in vitro* data is further supported by the *in-silico* prediction of a molecular interaction between EphA2 and CD44 such that CD44 may modulate canonical signaling of EphA2 via the LBD domain, while the FN-mediated interaction with CD44 may serve to stabilize the CD44-EphA2 complex near the cell surface and facilitate *Cn* entry.

**Fig 7.**
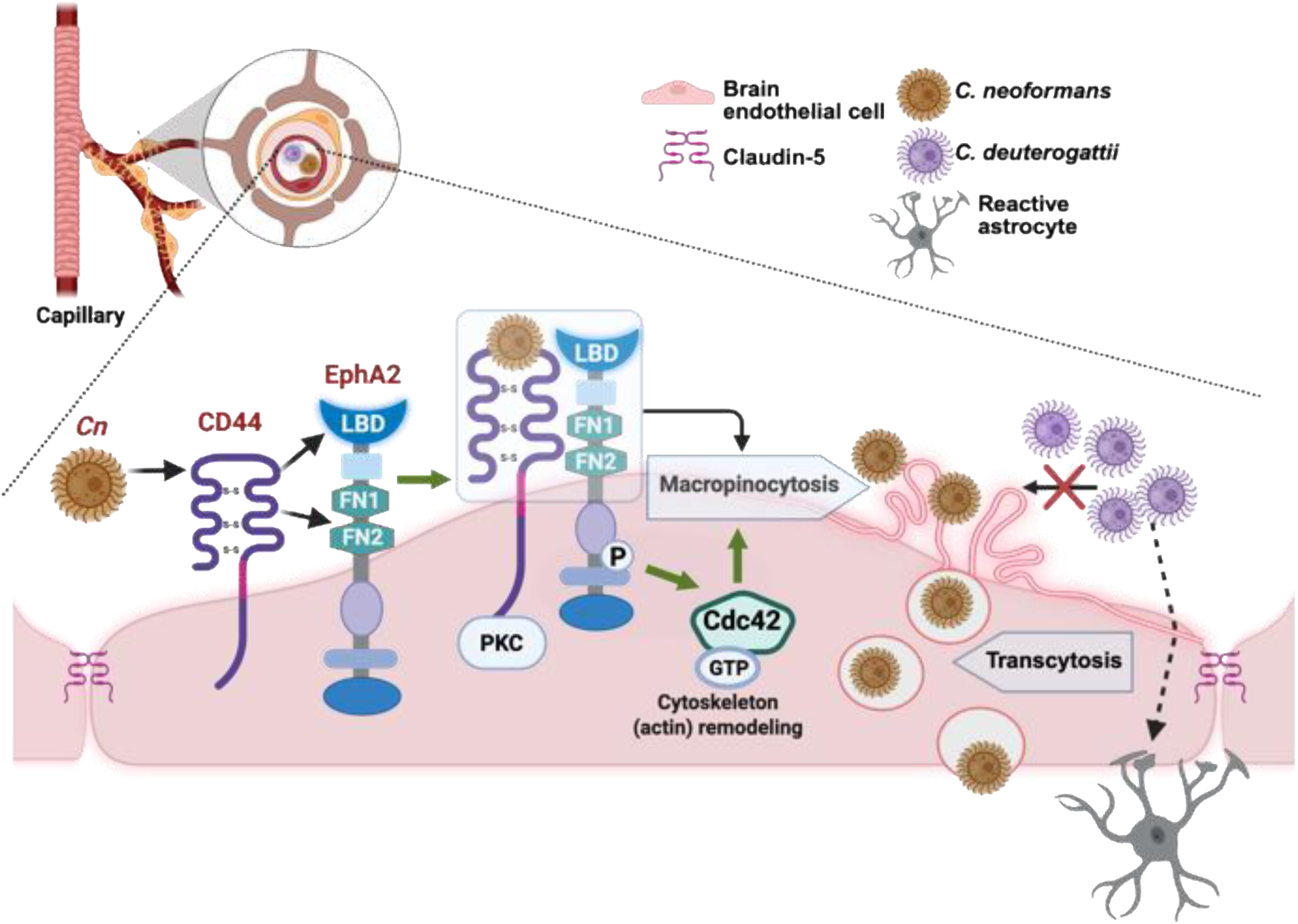
Conceptual model illustrating the molecular interactions that drive the transcytosis of *Cn* across the BBB. Upon binding to CD44, *Cn* promotes the association of CD44 with the ligand binding domain (LBD) and interactions involving both FN1 and FN2 domains, suggesting that CD44 can also engage EphA2 via a membrane-proximal site and/or the fibronectin 2 domain (FN2) (black arrows). The CD44-EphA2 molecular interaction leads to a PKC-dependent phosphorylation of EphA2, and the downstream activation of GTP-bound Cdc42 (green arrows). Active Cdc42 stimulates membrane ruffling by remodeling the cytoskeleton via actin polymerization. The ensuing macropinocytosis of *Cn* is a result of macropinosomes large enough to internalize and transport *Cn* across the brain endothelium. *Cd* does not cross the brain endothelium (crossed-out black arrow), but stimulates the expression of GFAP+ astrocytes by as yet an unknown mechanism (dashed arrow). Figure created in BioRender.com.

Consistent with macropinocytosis as the mode of entry, we found that amiloride treatment prevented the internalization of *Cn* by brain endothelial cells in both the BBB organoid model and the monoculture transwell BBB model. Amiloride is known to inhibit macropinocytosis by reducing endomembrane pH and preventing Cdc42 signaling, which is required for the actin-polymerization/remodeling needed to form the membrane ruffles and maturation of macropinosomes.^39,76^ Our results further demonstrate that brain invasion by *Cn* does not damage tight junctions or proceed through lytic death of the brain endothelium, at least initially.^22^

Despite their advantages, certain tradeoffs associated with BBB organoids must be noted. While the BBB organoid model recapitulates the cell-to-cell contact required for BBB function and barrier integrity, it still cannot recapitulate all the relevant *in vivo* conditions, such as the directional shear forces from blood flow. Also, the organoid core appears to be solid, unlike the “less compact” organization of pericytes and astrocytes observed in brain tissue, which likely impedes the penetration of *Cn* deep into the organoid interior. Our organoid model does not yet include neurons, oligodendrocytes, and microglia, which are likely to respond to internalized fungal cells and further impact the integrity of the BBB.^4,5^ However, we emphasize that our organoid model readily captures other important interactions between cells of the neurovascular unit and cryptococci; we found that astrocytes were especially activated upon exposure to *Cd*, and yet *Cd* had lower crossing rates than *Cn*, meaning that the effect of *Cd* on astrocytes is much stronger than *Cn* given that relatively few astrocytes were in direct contact with *Cd* as compared to *Cn*. This suggests that secreted factors from *Cd* may be responsible for the astrocyte activity observed in our studies.

Astrocyte activation can occur as a result of BBB dysfunction, with tight junctions breaking down completely and allowing macromolecules (e.g., albumin) to come into direct contact with astrocytes, eliciting inflammation.^80,81^ However, if this had occurred in our study, it did not occur to the extent that permitted *Cd* to infiltrate between endothelial cells. This implies an A2 type astrocytic response, in which astrocytes act to reinforce the BBB ^82–84^ and further suggests that *Cn* is entering the brain with the cooperation of brain endothelial cells, while *Cd* lacking this cooperation, elicits an astrocytic response, and is thwarted from crossing. However, further studies would be required to fully differentiate A1 and A2 astrocytes in the context of fungal neuroinvasion.^84^ That *Cd* has reduced BBB crossing relative to *Cn* in the BBB organoid model, is consistent with epidemiological studies demonstrating that *Cd* causes significantly fewer CNS infections compared to *Cn*.^85,86^ However, when *Cd* does infect the brain it often causes severe neurological complications likely due to a strong immune response that leads to inflammation.^87^

It is well known that capsule components of *Cn* downregulate the mammalian host immune response.^4,88^ The secreted *Cn* capsule component glucuronoxylomannan (GXM) in particular has been shown to influence the response of microglia to CNS-resident *Cn*, with cryptococcoma-proximal microglial cells adopting a non-phagocytic cell morphology near fungal cells producing a wild-type capsule.^4^ In the mouse model, GXM also reduces overall infiltration of microglia into cryptococcomas by directly reducing microglia motility, and skews the composition of circulating immune cells extravasating to sites of infection away from macrophages and B cells in favor of neutrophils and T-cells.^4^ The capsules of *Cn* and *Cd* differ in terms of capsule composition and average length of polysaccharides, with *Cd* expressing shorter GXM chains, and provoking a greater immune response in certain tissues,^49^ although *Cd* may have a certain advantage over *Cn* in immune evasion in the lung.^89^ Secreted capsule components that differ between *Cn* and *Cd* may therefore account for the differences in the astrocytic response observed in our BBB organoid model. Utilizing a human brain organoid model generated from human embryonic stem cells infected with *Cn* and *Cd* may provide further insight, as would expanding the human BBB organoid model to include additional cell types.^90^

### Methods Organoid model

Brain microvascular endothelial cells (HBMEC/D3 endothelial cells; also referred to as immortalized BMECs or iBMECs) were a gift from Babette Weksler.^40^ These iBMECs were maintained in collagen-coated flasks with Endothelial Growth Media (EGM, Lonza) containing 0.25% FBS, 1 ng/µL Fibroblast Growth Factor, and Penicillin/Streptomycin. Human primary astrocytes and brain vascular pericytes were purchased from ScienCell (Cat. #1800 and Cat. #1020 respectively, Carlsbad, California). Astrocytes and pericytes were maintained in poly-L-lysine coated flasks with their respective growth media (ScienCell, Carlsbad, California). In some experiments, primary human brain microvascular endothelial cells (Cat. #1000, ScienCell), maintained in the supplier-provided media (Cat. #1001, ScienCell) were used. Organoids were formed as previously described.^34^ Briefly, astrocytes and pericytes were cultured separately in 2D cultures before being trypsinized in 0.025% Trypsin-EDTA, seeded at 1×10^3^ per well each in low-adhesion polymer-coated 96-well plates (InSphero, Brunswick, Maine) inclined at 30 degrees to foster the formation of single organoids in each well for 48 h (37°C, 5% CO_2_). ECs were then split and seeded at 1×10^3^ cells per well on top of the astrocytes and pericytes.

### Baseline Permeability with FITC-dextran

Organoid permeability was established using low (FITC) versus high (FITC-dextran, 70 kDa) molecular weight fluid-phase fluorescent markers. FITC-dextran was diluted in EGM-2 media to a working concentration of 14.3 nM (10 µg/mL) and FITC to 500 nM (0.2 µg/mL) because it was empirically determined that these concentrations were approximately equivalent in relative fluorescence. Organoids were incubated in either FITC-dextran or FITC for 24 h, washed twice with warm PBS and once with media, then imaged on a confocal microscope at up to 100 µm depth in 8 µm slices. Average fluorescent signal in the FITC channel inside the spheroid at 100 µm was compared to assess permeability.

### Fixation and sectioning of organoids

Following treatment, organoids were removed from 96-well plates using flame-polished glass aspirators and placed in conical tubes containing immunofluorescence buffer (IF buffer: 0.15 M NaCl, 5 mM EDTA, 20 mM HEPES pH 7.5) and washed thrice, with intervals to settle to the bottom of tubes between washes. PBS was replaced with 4% PFA (pH 7.4 in PBS) and fixed overnight at 4°C with agitation. Organoids were then washed in IF buffer thrice before being placed in a 30% sucrose solution for cryoprotection, which was left to diffuse into organoids for 3 h at 4°C. DAPI was included in the cryoprotectant solution to aid visibility in a subsequent step. Flame-polished glass aspirators in combination with a stereoscope were used to transfer organoids to embedding molds and subsequently flash-frozen in a dry-ice ethanol bath and stored at −80°C. OCT blocks were sectioned at 15 µm in a cryostat under near-UV illumination in order to distinguish DAPI-stained organoids against the white OCT background. In experiments not requiring antibody staining, slides were rehydrated in IF buffer and sealed with aqueous mountant beneath coverslips.

### Live-cell membrane dye

An Invitrogen CellTracker fluorescent probe (Green, cat#: C7025) was used to stain pericytes prior to organoid formation to confirm the expected layering of cells in organoids. Staining procedure was done per the recommended manufacturer’s protocol. Briefly, the dye was dissolved in DMSO at 10 mM, then diluted in serum-free media to 15 µM. Cells were washed, trypsinized, pelleted at 200g for 3 min, and washed with warm PBS 3x. Cells were then incubated with dye-containing media for 30 min at 37°C in 5% CO_2_. Dyed cells were then washed twice with warm PBS before being combined with non-dyed cells as described above. Organoids were then fixed and sectioned as described previously before being imaged on a Leica TCS SP8 STED 3X confocal microscope with 50 nm laterally and 150 nm axially optical sectioning.

### Immunofluorescence

After sectioning, slides with spheroid-containing sections were brought up to room temperature over 30 min, then rehydrated in IF buffer for 1 min. Slides were then transferred to antigen retrieval solution (10 mM Sodium Citrate Buffer, 0.05% Tween 20, pH 6.0) for 30 min at 60°C, washed twice with IF buffer, then permeabilized and blocked for 2 h at room temperature in 1% BSA, 0.1 Triton X-100 in IF buffer. Antibodies [rabbit anti-Claudin5 (abcam ab15106-1003), 1:250, and mouse anti-GFAP (Invitrogen MA5-12023), 1:2000] were diluted in 1% BSA-IF buffer. Slides were incubated with primary antibodies at 4°C overnight, with rotation. Secondary antibodies (goat-anti rabbit TR ab6719, goat-anti mouse Cy5 ab6563, all used at 1:1000) were prepared in 1% BSA-IF buffer. Slides were washed five times in IF buffer, then incubated at room temperature with secondary antibodies for 2 h with gentle agitation. Sections were washed five times and nuclear stained with DAPI or Hoechst.

### TEM

Organoids were immersed in primary electron microscopy fixative (2.5% glutaraldehyde, 2% paraformaldehyde, 0.1 M sodium phosphate) for 3 h, rinsed with 0.1 M sodium phosphate, then fixed in secondary electron microscopy fixative (1% osmium tetroxide, 1.5 potassium ferrocyanide) for 1 h. Fixed organoids were washed thrice in cold deionized water, dehydrated in an ethanol series, then fixed in resin (Araldite 6005-Epon 812 mixture) overnight at 70°C. Resin blocks were then sectioned at 100 nm, transferred to copper grids, dried at 60°C for 30 min, and stained (4% uranyl acetate, 0.1 % lead citrate, 0.1 N sodium hydroxide). Imaging was performed with a FEI Talos L120C TEM 80kv.

### Generation of EphA2 knockout (KO) cell line

CRISPR/Cas9 technology was used to generate the HCMEC/D3 EphA2 KO cell line lacking expression of EphA2. Two sets of plasmids were used in the CRISPR/Cas9 method - the EphA2 CRISPR/Cas9 KO plasmids and the EphA2 HDR plasmids, both of which were ordered from Santa Cruz Biotechnology. The EphA2 CRISPR/Cas9 KO plasmids (Cat# sc-400535-KO-2) consist of a pool of three plasmids each encoding the Cas9 nuclease and a EphA2-specific 20 nt guide RNA (gRNA) designed for high knockout efficiency. Sequences of the gRNA were derived from the GeCKO (v2) library and direct the Cas9 protein to induce a site-specific double strand break (DBS) in the genomic DNA. Co-transfection with the EphA2 HDR plasmids (Cat# sc-400535-HDR-2) were carried out to select for cells containing a successful Cas9-induced DSB. The HDR plasmids consist of a pool of 2-3 plasmids, each containing a homology-directed DNA repair template corresponding to the cut sites generated by the EphA2 CRISPR/Cas9 KO plasmids. Each HDR plasmid contains two 800bp homology arms designed to specifically bind to the genomic DNA surrounding the corresponding Cas9-induced DBS site, leading to insertion a puromycin resistance gene to enable selection of stable knockout cells and insertion an RFP (Red Fluorescent Protein) gene to visually verify transfection. The puromycin resistance and RFP genes are flanked by two LoxP sites to allow for further processing by Cre recombinase (Cat# sc-418923) if desired. Transfection steps were carried out according to the manufacturer’s instructions. Briefly, freshly passaged HCMEC/D3 cells were first grown on a collagen-coated 6-well culture dish to about 50-80% confluence. The sub-confluent monolayer was then co-transfected for 48 h with 3 µg of EphA2 CRISPR/Cas9 KO plasmids and 3 µg of the EphA2 HDR plasmids in combination with 5-10 µl of the UltraCruz transfection reagent (Cat# sc-395739). After 48hours of incubation, transfected cells were grown in media containing 2.5 µg/mL of puromycin to select for successful recombination events in which a stretch of the HDR plasmid spanning the puromycin resistance gene and an RFP gene were integrated into the genomic DNA, leading to loss of transcription and expression of EphA2. Puromycin-resistant and RFP-positive cells were further purified and enriched using FACs (UC Davis flow cytometry core).

### Lentiviral Production and Transduction

To generate iBMECs overexpressing EphA2, lentiviral particles were first produced by transient transfection of HEK293T cells using a third-generation packaging system. Briefly, HEK293T cells were co-transfected with the transfer plasmid (pLenti-) and an optimized mix of packaging plasmids - pLP1, pLP2, and pLP/VSVG (ViraPower ^TM^ Lentiviral packaging mix, thermofisher scientific) using Lipofectamine 2000 (Thermo Fisher Scientific) according to the manufacturer’s instructions. Viral supernatants were collected 48 and 72 h post-transfection, filtered through a 0.45 µm PVDF filter, and stored at –80°C. iBMECs were transduced with viral supernatant at an MOI of 5 and supplemented with 8 µg/mL polybrene (Sigma-Aldrich). After 48 h, the medium was replaced, and cells were allowed to recover. Transduced cells were selected using blasticidin (1.5 µg/mL) for 3 days before analyzing expression via western blot analysis (anti-V5 tag antibody, ab27671, abcam inc. dilution 1:1000).

### BBB transcytosis assay

iBMECs were grown on collagen-coated transwells (Transwell-24 well plate, PET membrane, 8.0 µm pore size; Corning Inc., Lowell, MA, USA) as described previously.^40^ After differentiation of the cells, *Cn* at an MOI of 10 were added to the upper transwell chamber either in 1x EGM2 media to untreated cells (control treatment), in 1x EGM2 media to cells pretreated with 250 µm AML for 1 h (AML pretreatment) or in 1x EGM2 media with 250 µm AML to cells not pre-treated with AML (*Cn* and AML added together) to assess the effect of AML on the transcytosis of *Cn*. Cells were then incubated at 37°C, 5% CO_2_. At 12 h post-infection, the media in the lower chambers was collected and plated onto YPD agar plates to quantify the number of *Cn* that had crossed the iBMECs.

Organoid model (GFAP probing and quantification; *C. neoformans* vs *C. deuterogattii*; amiloride treatment assays)

Fungi (*Cn, Cd, Sc* strains, respectively - *Cryptococcus neoformans* H99, *Cryptococcus deuterogattii* VGII NIH-444, *Saccharomyces cerevisiae* W303) were grown overnight in 3 mL yeast peptone dextrose (YPD) liquid media at 30°C overnight. Second cultures were prepared 4 h prior to inoculation. Cultures were pelleted and washed in PBS 3x, diluted in PBS to 2×10 ^7^ cells/mL, then incubated in 500 µM CFDA-SE (10% DMSO) for 30 min at 37°C. The CFDA-SE-treated fungi were then washed and resuspended in cell culture media. Media over the organoids replaced with 100 µL of the fungal suspension, sufficient to completely bury organoids, which were returned to the incubator for 24-72 h, depending on the experiment. In experiments involving AML treatment, 250 µM AML or 0.25% DMSO (vehicle control) was added to the cell culture media along with fungi. Organoids were subsequently sectioned and imaged, as described above. BBB crossing was quantified in a semi-automated manner, with the edge of each organoid automatically delineated in ImageJ using a custom macro applied to the DAPI (cell nuclei) channel and redrawn on the FITC (fungi) channel (see supplementary information). This provided an unbiased boundary for manual counting of internalized versus adherent fungi. Fungal cells that had entered at least as far as the first layer of endothelial cells were considered as those that breached the organoid BBB.

### Tight junction protein localization in organoids

BBB organoids were created with either regular or EphA2 KO hCMEC/D3 cells, treated with *Cn* for 48 h, and sectioned as described above. Sections were probed with rabbit anti-Claudin5 (1:1000, ab15106, abcam) primary antibodies and goat anti-rabbit Cy5 (1:1000, ab6565, abcam) secondary antibodies. To create a metric of protein localization within organoids, pixel intensity in the Claudin5 (Cy5) channel across the diameter of organoid sections was measured using the Plot Profile tool in ImageJ. Values were normalized to the maximum value within each profile. The coefficient of variation for each normalized profile was then calculated in order to measure signal localization.

### 2D FITC-dextran uptake

iBMECs were grown on 4 well chambered slides (Nunc^TM^ Lab-Tek^TM^ ll Chamber slide^TM^ System, Thermo Fisher Scientific Inc., Waltham, MA) as previously described.^20,21^ Cells were either exposed to *Cn* (mCherry strain) at an MOI of 50 and 0.5 mg/mL of FITC-dextran or 0.5 mg/mL of FITC-dextran alone. Cells were incubated at 37°C, 5% CO_2_ for 1 h. Cells were immediately placed on ice, washed in 1x PBS (pH 7.4) and then washed in 0.1M NaCl solution and 0.1M Glycine solution (pH 3.00) 3x. After another 1x PBS wash, cells were fixed in 4% PFA for 10 min and prepared for confocal imaging (Leica, TCS SP8 STED 3x). For quantification of internalized dextran, an internalization index (dextran signal/area of the cell) was taken for 10 cells per treatment group using Image J software.

### Cdc42 Activation Assay

WT and EphA2 KO iBMECs were grown to confluency on 6-well plates as previously described^21^ (6-well plates, Corning Inc., Lowell, MA, USA). After differentiation, iBMECs were exposed to *Cn* at an MOI of 10 on ice and then incubated at 37°C, 5% CO_2_ for 1 h. To measure Cdc42 activation, the G-LISA activation kit (Kit # BK127 Cytoskeleton Inc., Denver, CO) were used per manufacturer’s recommendations. This assay used Cdc42-GTP binding proteins linked to the wells of a 96-well plate. Active, GTP-bound Cdc42 in cell lysates bound to the wells while inactive GDP-bound forms were removed during wash steps. Bound GTP Cdc42 were detected by incubation with a Cdc42 specific antibody followed by a secondary antibody conjugated to HRP and a detection reagent. The signal was quantified by measuring absorbance at 490 nm using a microplate reader (Spectramax M5, Molecular Devices Inc., San Jose, CA).

### Western Blot Analysis

Strains of *C. neoformans* (*Cn* WT and *cps1Δ*) were lysed with RIPA buffer (10X RiPA Buffer abcam Inc.) supplemented with a protease inhibitor (cOmplete mini, EDTA free cocktail protease inhibitor, Roche Inc.) and a phosphatase inhibitor (Roche PhosStop phosphatase inhibitor, Roche Inc.). The protein concentrations were measured by Bradford Assay (Quick Start^TM^ Bradford Protein Assay; Bio-Rad Laboratories Inc.), and the samples were denatured and reduced in Laemmli buffer (Premixed 4X Laemmli protein sample Buffer; Bio-Rad Laboratories Inc.) and 2-mercaptoethanol (Bio-Rad laboratories Inc.), followed by boiling at 100°C for 5 min. SDS-PAGE electrophoresis was performed with 10% polyacrylamide gel at 80 V for 3 h (Mini-PROTEAN Tetra Cell; Bio-Rad laboratories Inc.). The proteins on the SDS PAGE gel were transferred to a nitrocellulose membrane (Nitrocellulose/ Filter paper sandwiches 0.45µm, Bio-Rad Inc.) using the wet transfer method at 4°C (Mini Trans-Blot Cell; Bio-Rad Inc.), 30 V overnight for analyzing the EphA2 protein and 100 V for 2 h for analyzing the Cdc42 protein. The nitrocellulose membrane was stained with Ponceau Red (PONCEAU S High Purity; VWR Inc. 0.1% wt/vol with glacial acetic acid 5% vol/vol) to visualize polypeptide bands and then blocked with 5% milk (Blotting-Grade Blocker; Bio-Rad Inc.) for 1 h at room temperature. Membranes were then incubated with primary antibody (1:1000 dilution of Rb anti-Human EphA2; Cell signaling technology Inc. D4A2) (1:800 dilution of Ms anti-human Cdc42; cytoskeleton Inc. ACD04) at 4°C overnight. The membrane was washed several times with TBST buffer and then incubated in secondary antibody (1:10,000 dilution of Goat anti-Mouse IgG H&L HRP ab6789 and Goat Anti-Rabbit IgG H&L HRPab6721; Abcam Inc., Cambridge, MA, USA) at room temperature for 1 h. After washing with TBST, the chemiluminescent substrate solution for HRP (Supersignal West Pico Chemiluminescent substrate; Pierce Biotechnology, Rockford, IL, USA) was added to the membranes and X-Ray films (HyBlot ES Autoradiography Film; Denville Scientific Inc., Metuchen, NJ, USA) were used to detect the expression of EphA2 and Cdc42 proteins.

### EphA2-CD44 Proximity Ligation Assay (PLA)

Brain endothelial cells (BECs) were seeded at 50% confluence in collagen-coated LabTek II chamber slides in EGM2 media containing 0.25% FBS and 1 ng/µL FGF (1X media). To promote the BBB phenotype, after the endothelial monolayer had reached confluency, the concentration of growth factors in media was reduced by half for 24 h, then reduced by half again for another 24 h before introduction of cryptococci (*Cn* or *Cd*). Cryptococci were grown at 30°C overnight in YPD, then secondary cultures were prepared from overnight cultures and grown 4 h prior to experiments to ensure that fungi were in logarithmic phase growth. The secondary cultures were washed thrice in PBS, then diluted to deliver 3.5×10^5^ cryptococci per 0.7 cm^2^ chamber (MOI of 5). Cryptococci and BECs were co-incubated at 37°C, 5% CO_2_ for 90 minutes. Control chambers that did not receive fungi were prepared as well. Cells were then washed with warm PBS and fixed with 4% PFA for 10 min at room temperature. A proximity-ligation assay (PLA) using the Duolink system (Sigma-Aldritch) was conducted following the supplier’s instructions (protocol source: https://www.sigmaaldrich.com/deepweb/assets/sigmaaldrich/marketing/global/documents/154/8 82/introduction-duolink-pla-4383-ms.pdf). Briefly, after fixation, cells were blocked with Duolink blocking solution for 1 h at 37°C in a humidified chamber. Slides were then incubated with mouse anti-CD44 (Abcam ab254530) and rabbit anti-EphA2 (Abcam ab185156), diluted 1:500 and 1:50, respectively, in Duolink antibody diluent (Sigma-Aldrich), and incubated at 4°C overnight with agitation. Slides were washed twice with PBS before being incubated with anti-mouse PLUS and anti-rabbit MINUS probes diluted 1:5 in DuoLink antibody diluent and incubated for 1 h in a humidified chamber. Ligase (1:40 in ligation buffer) was then added to ligate PLUS and MINUS probes (up to 40 nm separation, per manufacturer specifications) and incubated for 30 min at 37°C. Amplification buffer containing the red detection reagent and polymerase (1:80) were incubated with slides for 100 min at 37°C. Control slides that did not receive primary antibodies were processed in parallel. After cover slips were added, slides were imaged using a fluorescence microscope. Quantification of fluorescence was performed in ImageJ.

### Molecular Dynamics Simulations

We utilized Alphafold3^52^ (ref) to model the interactions of extracellular region (ECR) of EphA2 (A24-V537) with CD44 (Q21-S180). To analyze the protein-protein interface between EphA2-CD44, we started our md simulations with two systems: first one with the LBD (K27-L205) of EphA2 with CD44 and the second one with the FN domains (C325-E530) of EphA2 with CD44. As a control system we used the LBD of EphA2 with EphrinA5 from the crystal structure (PDB: 2X11). The FN:CD44 system and the LBD:CD44/ EphrinA5 systems were encased in a box with dimensions measuring 110 × 110 × 110 Å^3^ and 90 × 90 × 90 Å^3^ respectively. The system was solvated using the TIP3P water model. To ensure neutrality, Na+ ions were added and also 150mM of NaCl was introduced using the CHARMM-GUI interface.^91^ Molecular dynamics simulations were performed using GROMACS 2023.3^92^ with the CHARMM36m force field.^93^ Production MD simulations were carried out in GROMACS using the leap-frog integrator with a 2 fs timestep and a total length of 200 ns. Non-bonded interactions were treated with the Verlet cutoff scheme, a 1.2 nm cutoff (force-switch for van der Waals), and PME for electrostatics. Temperature was maintained at 303 K using the velocity-rescale thermostat, and pressure at 1 bar with the C-rescale barostat. All bonds involving hydrogens were constrained with LINCS. A total of three replica simulations were conducted for each system. Trajectory analysis was conducted using the integrated modules within GROMACS. Subsequently, the data was visualized and plotted using Origin.

### Statistical analysis

Data were analyzed for statistical significance with either GraphPad Prism 10 software or R v4.4.2. Statistical analysis of two data sets was performed using unpaired, two-sided parametric t-tests with Welch’s correction. Data sets greater than two were analyzed by one-way analysis of variance (ANOVA) with Tukey’s multiple comparisons test. (ns = not significant, \**p*<0.05, \*\**p*<0.01, \*\*\**p*<0.001, \*\*\*\**p*<0.0001).

## Conflict of interest

The authors have declared that no conflict of interest exists.

---

**Supplementary Fig. 1.**
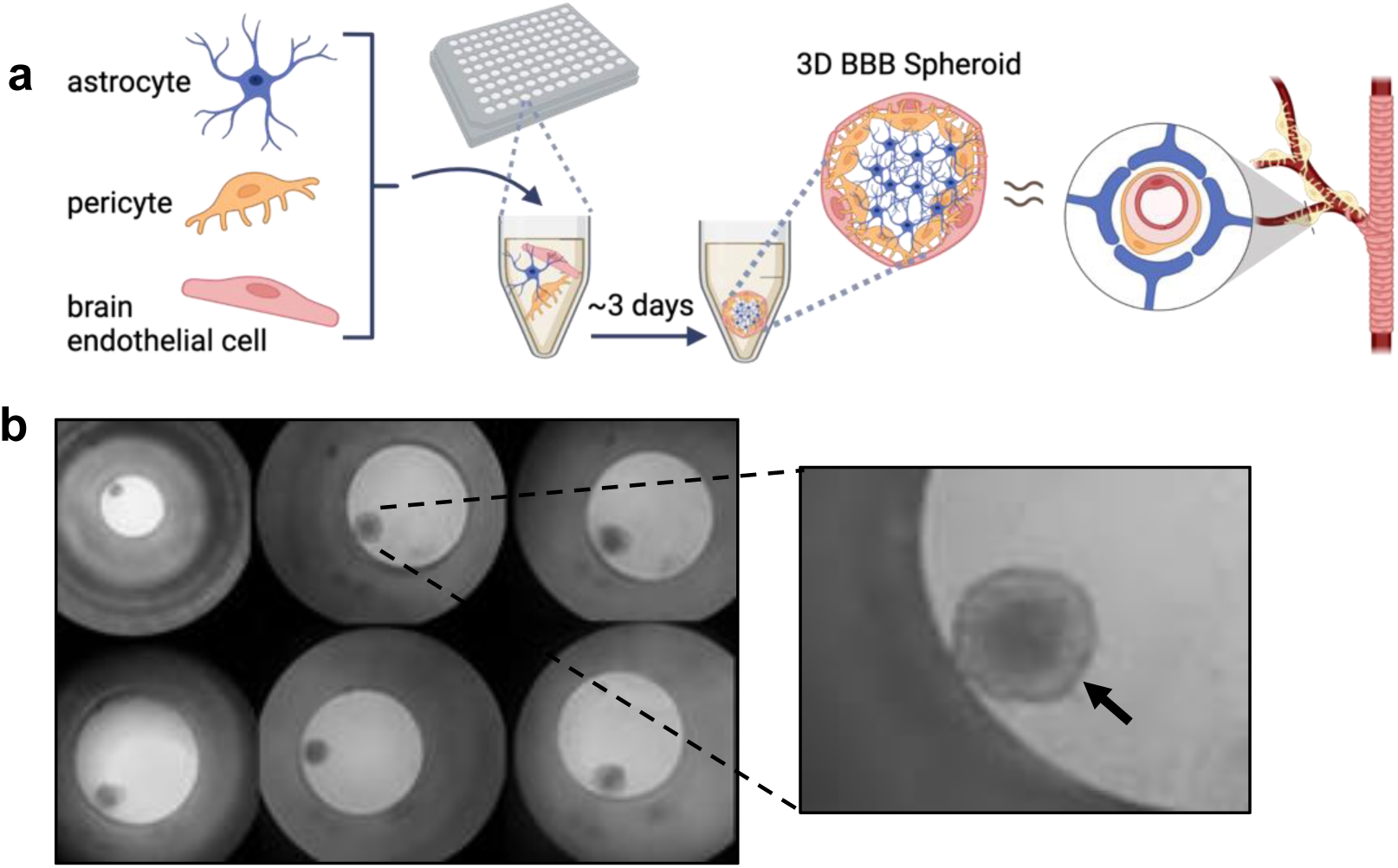
**Schematics and images of 3D BBB organoid formation**. **a** Organoids were formed from either primary human BMECs (passage < 6) or immortalized HCMEC/D3 cells (iBMECs, passage < 35), human primary pericytes (passage 4), and human astrocytes (passage 4). Primary cells were purchased from ScienCell (Carlsbad, California, USA). The organoids were formed by combining the three cell types in a 1:1:1 ratio, each at a concentration of 1×10^4^ cells/mL. The combined cells were grown in 100 µL media in a 96-well plate. **b** Brightfield view of organoids formed at the bottom of flat-bottom 96-well plates with a transparent non-stick polymer coating. Organoids were relatively homogenous in size and shape.

**Supplementary Fig. 2.**
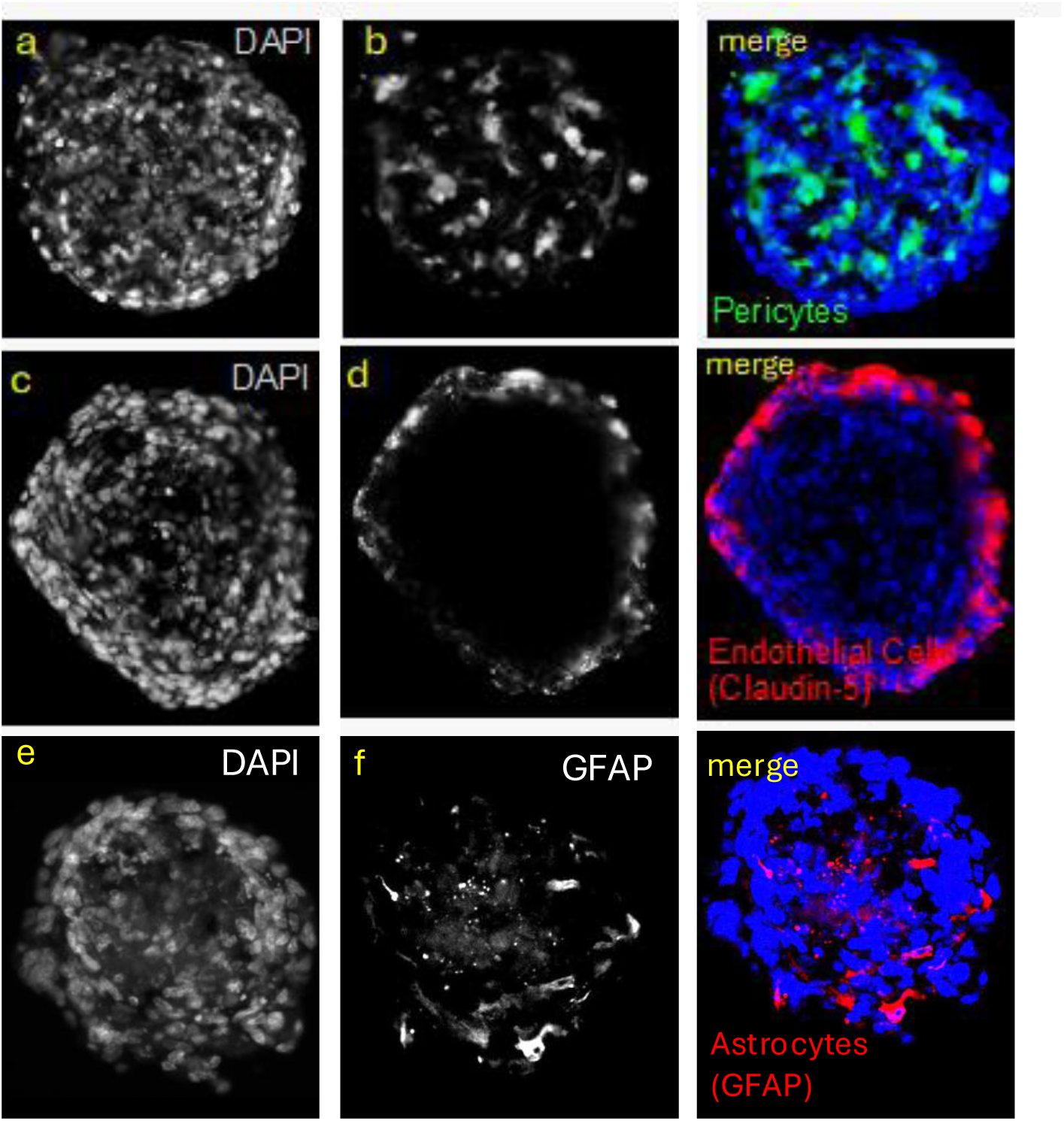
Organization of neurovascular cell types in human BBB organoids. First column **a**, **c**, **e** shows DAPI counterstaining. Second column **b**, **d**, **f** shows cell-type specific staining. Third column contains merged images from column 1 and column 2. Top row: pericytes were labeled in culture with CMFDA (CellTracker Green, Invitrogen) and combined with endothelial cells and astrocytes to form organoids. **a** DAPI, **b** pericytes, merged image to the right. Middle row: Organoids were probed with **d** rabbit anti-Claudin-5 (1:250) and goat - anti-rabbit TexasRed, merged image (middle row, right) with DAPI **c**. Bottom row: Astrocytes were visualized with GFAP (Invitrogen MA5-12023, 1:2000) **f** with DAPI counterstaining **e**, with merged images to the right. All images were acquired from 15 µm thick cryosections of PFA-fixed sections made from 1-week old organoids.

**Supplementary Fig. 3.**
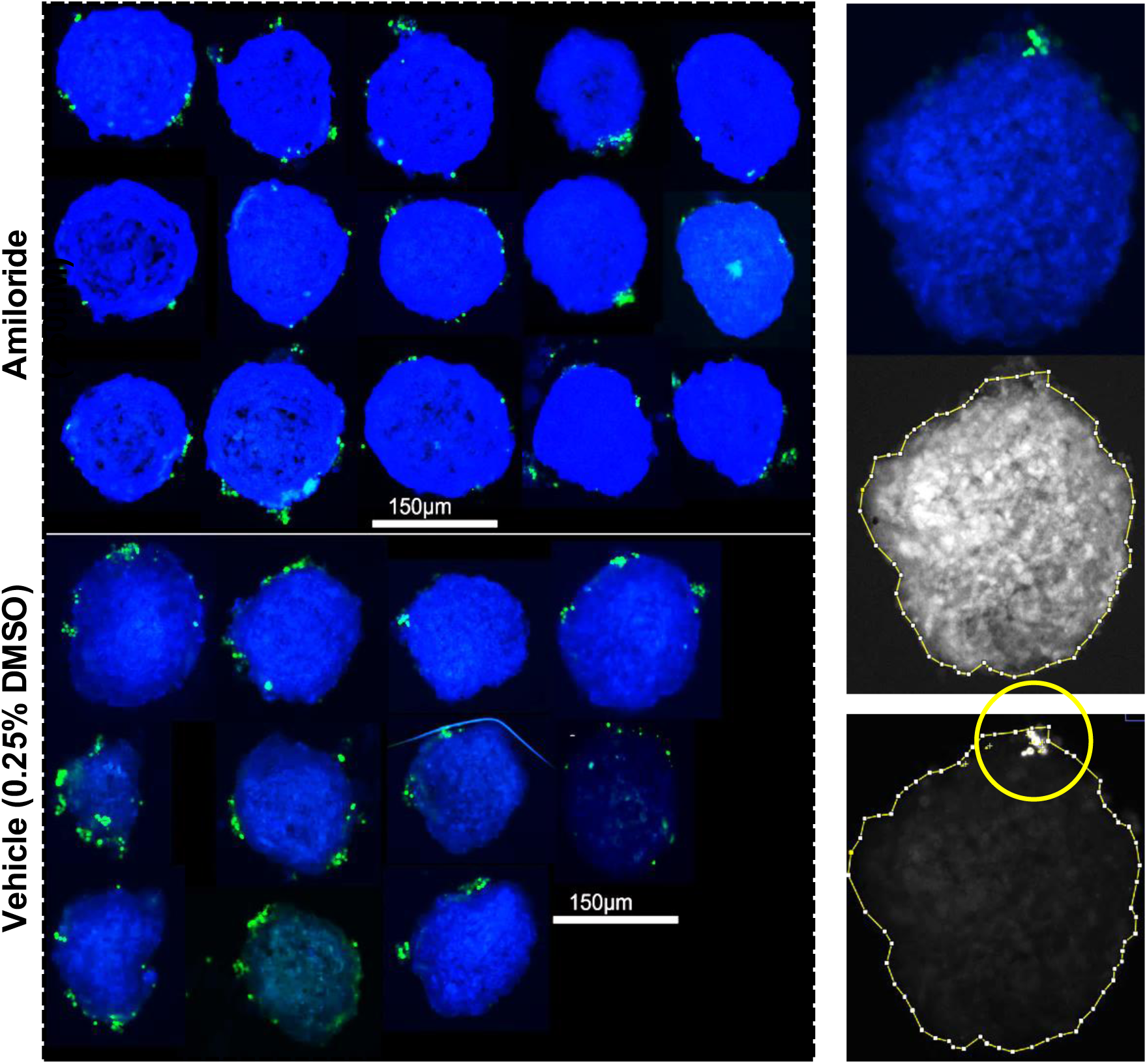
Organoid-based assays to examine the impact of Amiloride treatment on *Cn* BBB-internalization/crossing. Representative images showing organoids (blue, DAPI) incubated for 24 h with *Cn* (green, CFSE) in media containing either 250 µM amiloride (AML) (top-left) or an equivalent concentration of vehicle control (bottom left). Representative images from the analysis pipeline, with the “input” merged image at top right, border demarked around organoids in the DAPI channel right middle panel, and this border applied in the CFSE channel to quantify crossing/internalization of fungal cells (bottom right panel, yellow circle).

**Supplementary Fig. 4.**
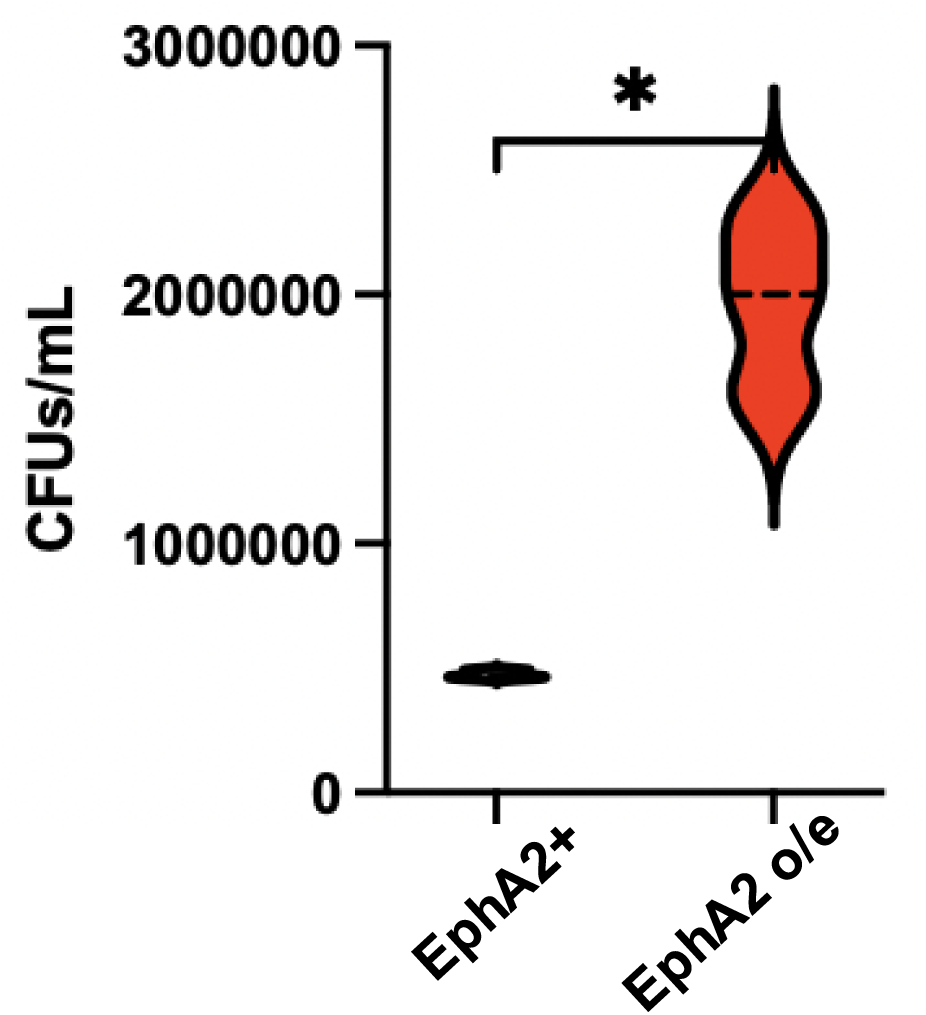
Overexpression of EphA2 in brain endothelial cells increases migration of *Cn* across the brain endothelium in an *in vitro* transwell model of the BBB. Transcytosis assay shows a significant increase in the movement of *Cn* across endothelial cells when brain endothelial cells (iBMECs) overexpress EphA2. EphA2+ iBMECs (wild type) and iBMECs transduced with EphA2 *via* lentivirus were exposed to *Cn* (MOI of 10) that was added to the apical side of the transwell. Following 12 h exposure, fungal cells were collected from the abluminal side of each transwell (well below transwell) and plated onto agar plates for CFU (colony forming units) determination. Each data point represents the fungal count from an individual well (n=3). Statistical analysis was done using a two-tailed, unpaired t-test with Welch correction (*p* = 0.0178, t= 7.358, df= 2.008)

**Supplementary Fig. 5.**
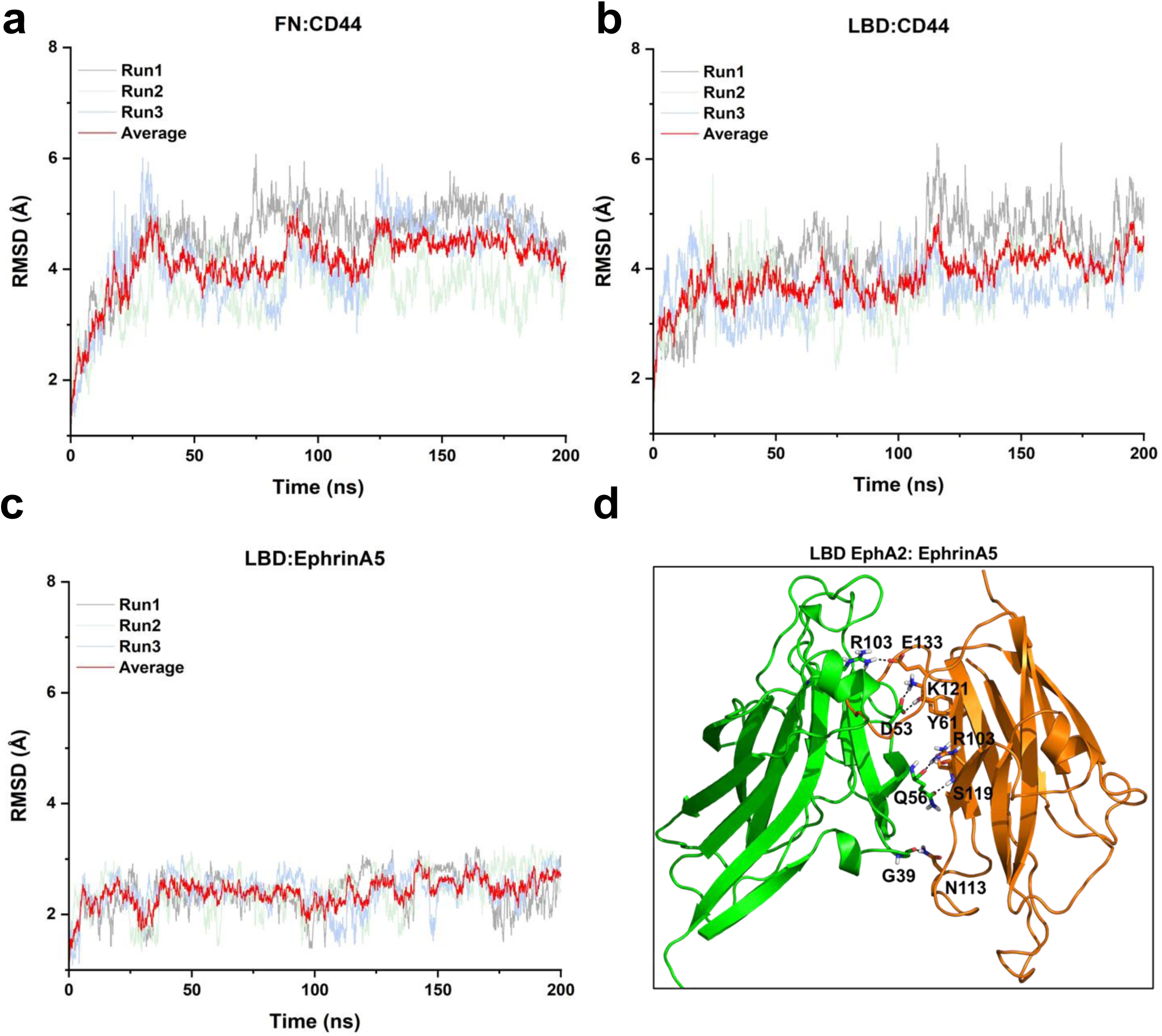
Structural stability and binding conformations of EphA2 complexes. a–c. Backbone RMSD profiles of the FN: CD44, LBD: CD44, and LBD: ephrinA5 complexes over the course of molecular dynamics simulations. Data from all replica trajectories as well as averaged RMSD values are shown. **d** Representative binding conformation of the LBD: ephrinA5 complex obtained from clustering of all replica trajectories. EphA2 is depicted in green and ephrinA5 in orange cartoon representation. For clarity, only the major interfacial hydrogen bonds are displayed.

**Supplementary Fig. 6.**
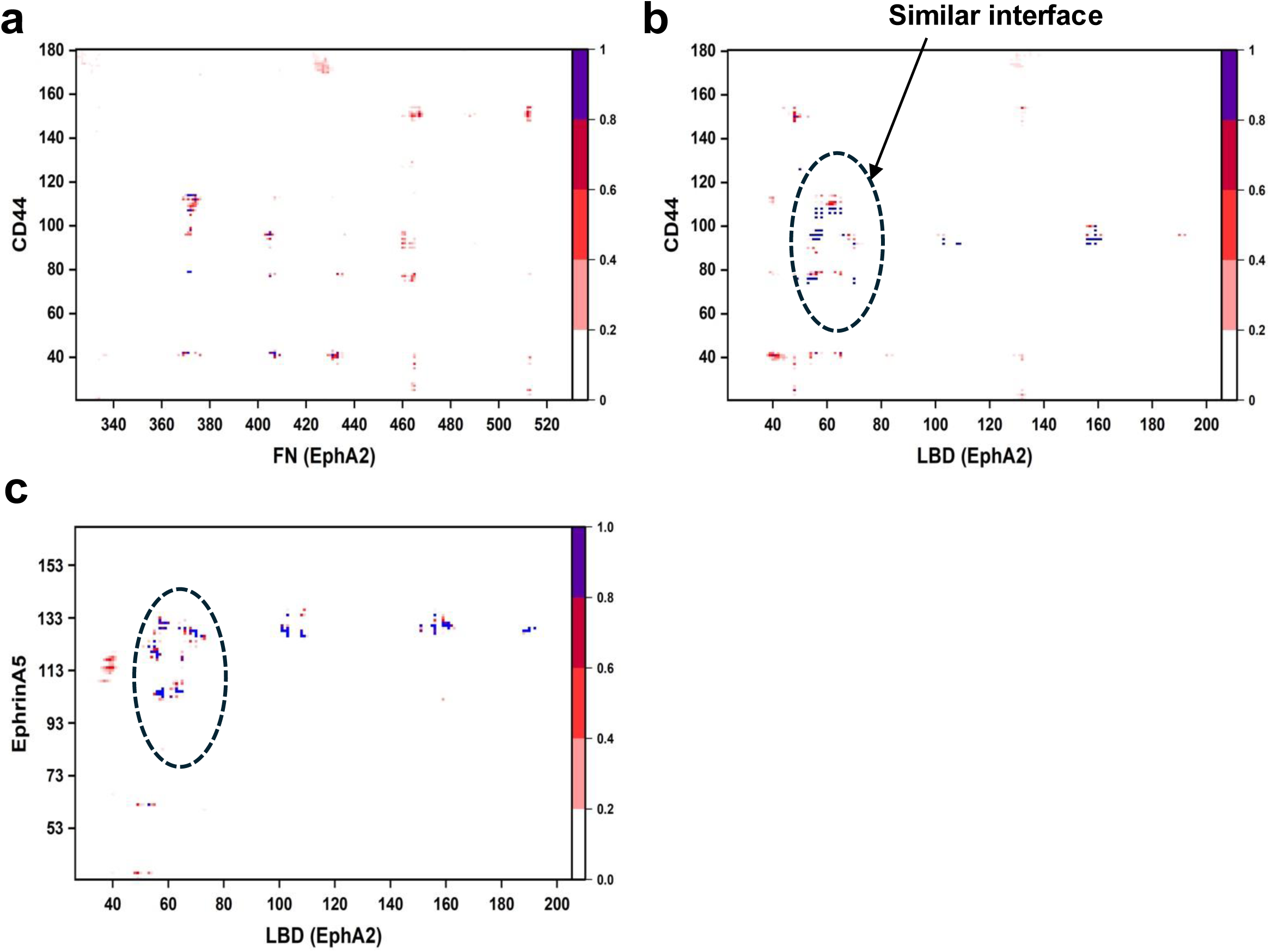
Intermolecular contact interfaces of EphA2 complexes. (**a–c**) Contact maps of the FN: CD44, LBD: CD44, and LBD: ephrinA5 complexes obtained from molecular dynamics simulations. Contacts were calculated using a 5 Å cutoff. The color scale (white to red to blue) represents the fractional occupancy of each contact, ranging from 0 (no contact) to 1 (persistent contact). Data from all replica trajectories were included in the analysis. Notably, CD44 and ephrinA5 engage overlapping interface regions within the EphA2 LBD.

**Supplementary Table 1.** Antibodies used in the current study.

| **Name** | **Type** | **Company** | **Catalog #** | **Use** |
| --- | --- | --- | --- | --- |
| mouse anti-CD44 | primary | abcam | ab254530 | PLA |
| rabbit anti-EphA2 | primary | abcam | ab185156 | PLA |
| mouse anti-Cdc42 | primary | cytoskeleton | ACD04 | WB |
| rabbit anti-EphA2 | primary | Cell signaling | D4A2 | WB |
| goat anti-mouse IgG<br>H&L HRP | secondary | abcam | ab6789 | WB |
| goat anti-rabbit IgG<br>H&L HRP | secondary | abcam | ab6721 | WB |
| rabbit anti-Claudin5 | primary | abcam | ab15106 | IF |
| goat anti-rabbit Cy5 | secondary | abcam | ab6564 | IF |
| mouse anti-GFAP | primary | Invitrogen | MA5-12023 | IF |
| goat anti-rabbit<br>TexasRed | secondary | abcam | ab6719 | IF |
| goat anti-mouse Cy5 | secondary | abcam | ab6563 | IF |

## Notes

### Competing Interest Statement

The authors have declared no competing interest.

### Funding

- National Institutes of Health: NS110800

